# Distinct progenitor populations mediate regeneration in the zebrafish lateral line

**DOI:** 10.1101/482737

**Authors:** Eric D. Thomas, David W. Raible

**Affiliations:** Department of Biological Structure, University of Washington, 1959 NE Pacific St, Box 357420, Seattle, WA 98195, USA; Graduate Program in Neuroscience, University of Washington, 1959 NE Pacific St, Box 357420, Seattle, WA 98195, USA; Virginia Merrill Bloedel Hearing Research Center, University of Washington, 1959 NE Pacific St, Box 357420, Seattle, WA 98195, USA

## Abstract

Mechanosensory hair cells of the zebrafish lateral line regenerate rapidly following damage. These renewed hair cells arise from the proliferation of surrounding support cells, which undergo symmetric division to produce two hair cell daughters. Given the continued regenerative capacity of the lateral line, support cells presumably have the ability to replenish themselves. Utilizing novel transgenic lines, we identified support cell populations with distinct progenitor identities. These populations show differences in their ability to generate new hair cells during homeostasis and regeneration. Targeted ablation of support cells reduced the number of regenerated hair cells. Furthermore, progenitors regenerated after targeted support cell ablation in the absence of hair cell damage. We also determined that distinct support cell populations are independently regulated by Notch signaling. The existence of independent progenitor populations could provide flexibility for the continued generation of new hair cells under a variety of conditions throughout the life of the animal.

## INTRODUCTION

The regenerative potential of a given tissue is dependent on the availability of progenitor cells that are able to functionally replace lost or damaged cells within that tissue. For instance, bulge cells in the hair follicle can repair the surrounding epidermis (Rompolas and Greco 2014; Hsu, Li, and Fuchs 2014), new intestinal epithelial cells arise from crypt cells (Santos et al. 2018; Yousefi, Li, and Lengner 2017), and horizontal and globose basal cells can regenerate cells in the olfactory epithelium (Choi and Goldstein 2018; Schwob et al. 2017). Depletion of these progenitors can severely diminish the regenerative capacity of the tissue, and tissues that lack a progenitor pool altogether are unable to regenerate. To gain further insight into how different tissues regenerate, a greater understanding of the mechanisms that define and regulate progenitor function are needed.

The zebrafish lateral line system has long been recognized as an excellent model for studying regeneration. The sensory organ of the lateral line, the neuromast, is comprised of mechanosensory hair cells organized on the surface of the head and body (Thomas et al. 2015). Lateral line hair cells regenerate rapidly following damage, with the system returning to quiescence after regeneration is complete (Harris et al. 2003; Hernandez et al., 2007; Ma et al., 2008). The surrounding nonsensory support cells serve as progenitors for new hair cells. This replenishment is proliferation-dependent and occurs symmetrically, with each progenitor dividing to give rise to two daughter hair cells (Wibowo et al. 2011; Mackenzie and Raible 2012; López-Schier and Hudspeth 2006; Romero-Carvajal et al. 2015). Three key observations of support cell behavior during regeneration suggest that different support cell populations may be differentially regulated in response to regeneration. First, the support cell proliferation that follows hair cell death occurs mainly in the dorsal and ventral compartments of the neuromast (Romero-Carvajal et al. 2015), indicating that progenitor identity is spatially regulated. The most peripheral support cells, often called mantle cells, do not proliferate in response to hair cell damage (Ma, Rubel, and Raible 2008; Romero-Carvajal et al. 2015). Second, the regenerative capacity of the neuromast is not diminished over multiple regenerations (Cruz et al. 2015; Pinto-Teixeira et al. 2015), indicating that progenitor cells must also be replaced in addition to hair cells. Finally, in addition to regeneration in response to acute damage, lateral line hair cells undergo turnover and replacement under homeostatic conditions (Cruz et al. 2015; Williams and Holder 2000). However, it remains unknown whether there are distinct support cell populations within the neuromast (e.g. hair cell progenitors and those that replenish progenitors), as well as how progenitor populations are regulated.

In this study, we have used CRISPR to generate novel transgenic lines in which distinct, spatially segregated populations of support cells are labeled *in vivo*. Fate mapping studies using these lines show that these populations are functionally distinct with respect to their ability to contribute new hair cells during homeostasis and to generate hair cells after damage. We also show that targeted ablation of one of these populations significantly reduces hair cell regeneration. Other fate mapping studies show that these support cell populations can replenish each other in the absence of hair cell damage. Finally, we show that Notch signaling differentially regulates these populations. These results demonstrate that there are a number of distinct progenitor populations within lateral line neuromasts that are independently regulated, providing flexibility for hair cell replacement under a variety of circumstances.

## RESULTS

### Hair Cell Progenitors are Replenished via Proliferation of Other Support Cells

Previous studies have shown that the majority of support cell proliferation occurs during the first twenty-four hours following hair cell death (Ma, Rubel, and Raible 2008). We replicated this finding by administering a pulse of F-ara-EdU (EdU), which has been shown to be far less toxic than BrdU (Neef and Luedtke 2011). The EdU pulse was administered for twenty-four hours following neomycin-induced hair cell ablation at 5 days post fertilization (5 dpf) and neuromasts were imaged at seventy-two hours post treatment (72 hpt), the time at which regeneration is nearly complete (Fig. 1A). In neomycin-treated larvae, roughly 78% of regenerated hair cells were EdU-positive, compared to 6% in mock-treated larvae (Fig. 1B-C; p< 0.0001). We noticed that at the same timepoint that 28% of EdU-positive cells remained support cells (Fig. 1E,1B arrowheads). We hypothesized that these cells may represent hair cell progenitors that had been replaced via proliferation. If so, then these EdU-positive cells should have the capacity to generate a new round of hair cells after subsequent damage. In order to test this, we subjected larvae to two rounds of hair cell ablation and regeneration. EdU was administered for 24h following the first ablation, and BrdU was administered for the same duration following the second ablation (Fig. 1F). We observed hair cells after the second regeneration that were both EdU- and BrdU-positive (Fig. 1G-J, arrowheads), indicating that support cells that divide after the first ablation can in fact serve as hair cell progenitors after subsequent damage. However, we also observed double-positive cells that remained support cells (Fig. 1K-N, arrowheads), as well as support cells that were only labeled by EdU (Fig. 1G-N, asterisks). These observations indicate that support cells that divided after the first round of damage do not always serve as hair cell progenitors (or even as progenitors at all). Altogether, these data provide evidence of proliferation-mediated replenishment of hair cell progenitors by support cells in the neuromast, but also suggest that new hair cells arise from a pool of progenitors that are not strictly defined by their proliferation history.

**Figure 1.**
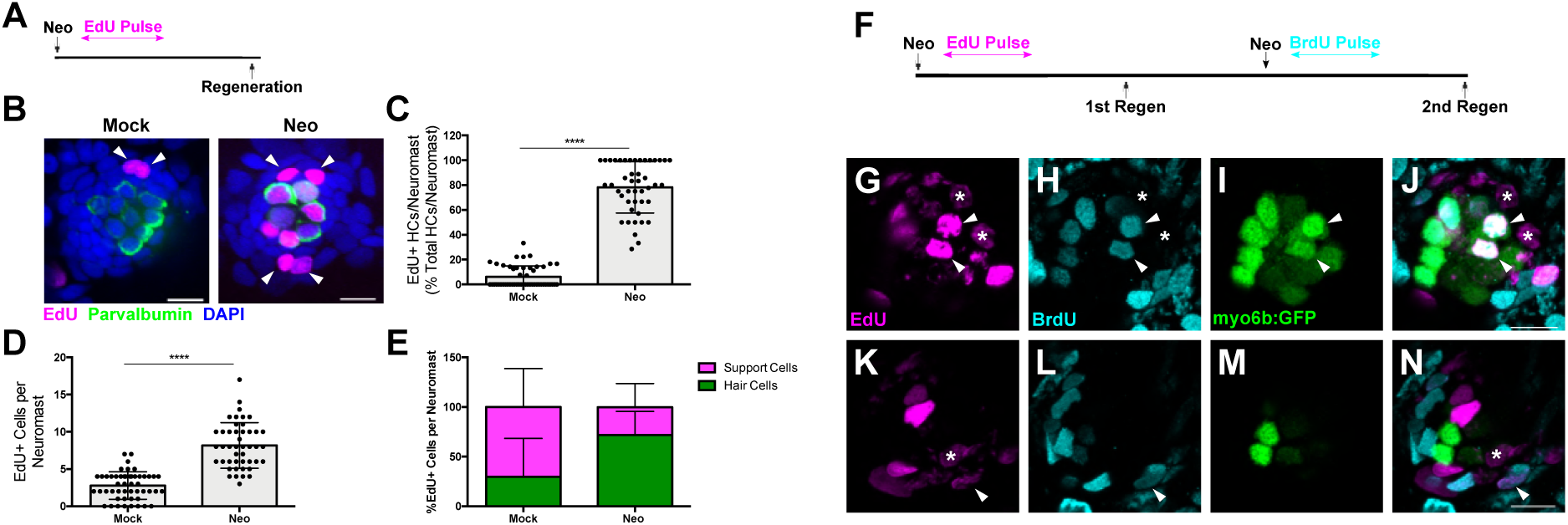
Hair cell progenitors are replenished via proliferation of other support cells. **(A, F)** Timelines of single-ablation (A) and double-ablation (F) proliferation experiments. (**B**) Maximum projections of mock-(Mock) and neomycin-treated (Neo) neuromasts. EdU-positive cells are shown in magenta, anti-Parvalbumin-stained hair cells are shown in green, and DAPI-stained nuclei are shown in blue. Arrowheads indicate EdU-positive support cells. Scale bar = 10 μm. **(C)** Percentage of hair cells per neuromast labeled by EdU. Mock: 6.11 ± 8.69, n = 50 neuromasts; Neo: 78.24 ± 20.69, n = 45 neuromasts; mean ± SD; Mann Whitney U test, p < 0.0001. **(D)** Total EdU-positive cells per neuromast. Mock: 2.78 ± 1.84, n = 50 neuromasts; Neo: 8.18 ± 3.07, n = 45 neuromasts; mean ± SD; Mann Whitney U test, p < 0.0001. **(E)** Percentage of EdU-positive cells per neuromast that are either hair cells or support cells. Mock: 29.73% hair cells, 70.27% support cells, n = 50 neuromasts; Neo: 72.02% hair cells, 27.98% support cells, n = 45 neuromasts; mean ± SD. **(G-N)** Individual slices of a neuromast following two regenerations at two different planes: apical hair cell layer (G-J) and basal support cell layer (K-N). EdU (visualized by a Click-iT reaction) is labeled in magenta, BrdU (anti-BrdU) is labeled in cyan, and myo6b:GFP hair cells are labeled in green. Arrowheads indicate EdU/BrdU-positive hair cells, and asterisks indicate EdU-positive support cells. Scale bar = 10 μm.

### Different Progenitor Identities Among Distinct Support Cell Populations

We next sought to determine whether hair cell progenitors could be defined via gene expression. To this end, we employed CRISPR-mediated transgenesis (Kimura et al. 2014; Ota et al. 2016) to target genes that label positionally-defined subsets of support cells *in vivo*. These efforts were part of a broader insertional screen to be described elsewhere. We targeted the expression of a nuclear-localized form of the protein Eos (nlsEos) to a variety of genetic loci, and identified three genes (*sfrp1a, tnfsf10l3*, and *sost*) which have markedly different expression patterns within support cells: *sfrp1a* is restricted to the most peripheral support cells (Peripheral cells; Fig. 2A); *tnfsf10l3* is more broadly expressed throughout the periphery but is enriched in anteroposterior support cells (AP cells; Fig. 2C); and *sost* is limited to the dorsal and ventral support cells (DV cells; Fig. 2E). We generated stable transgenic lines for all three loci: Tg[*sfrp1a*:nlsEos]^w217^; Tg[*tnfsf10l3*:nlsEos]^w218^; and Tg[*sost*:nlsEos]^w215^ (hereafter known as *sfrp1a*:nlsEos, *tnfsf10l3*:nlsEos, and *sost*:nlsEos, respectively). Eos is a photoconvertible protein that switches from green to red fluorescence (shown in magenta throughout this paper) after exposure to UV light (Wiedenmann et al. 2004). The converted protein is stable for months. Its nuclear localization presumably protects it from degradative elements in the cytoplasm, allowing for a more permanent label than a standard fluorescent reporter (Cruz et al. 2015; McMenamin et al. 2014). We could thus chase this label from support cell to hair cell if these cells serve as hair cell progenitors, as hair cells that derived from these support cells would have converted Eos in their nuclei. To ensure that these genes were not actually expressed in hair cells, we also generated GFP lines for each gene. We did not observe GFP labeling in hair cells in stable lines (Fig. 2 – figure supplement 1).

**Figure 2.**
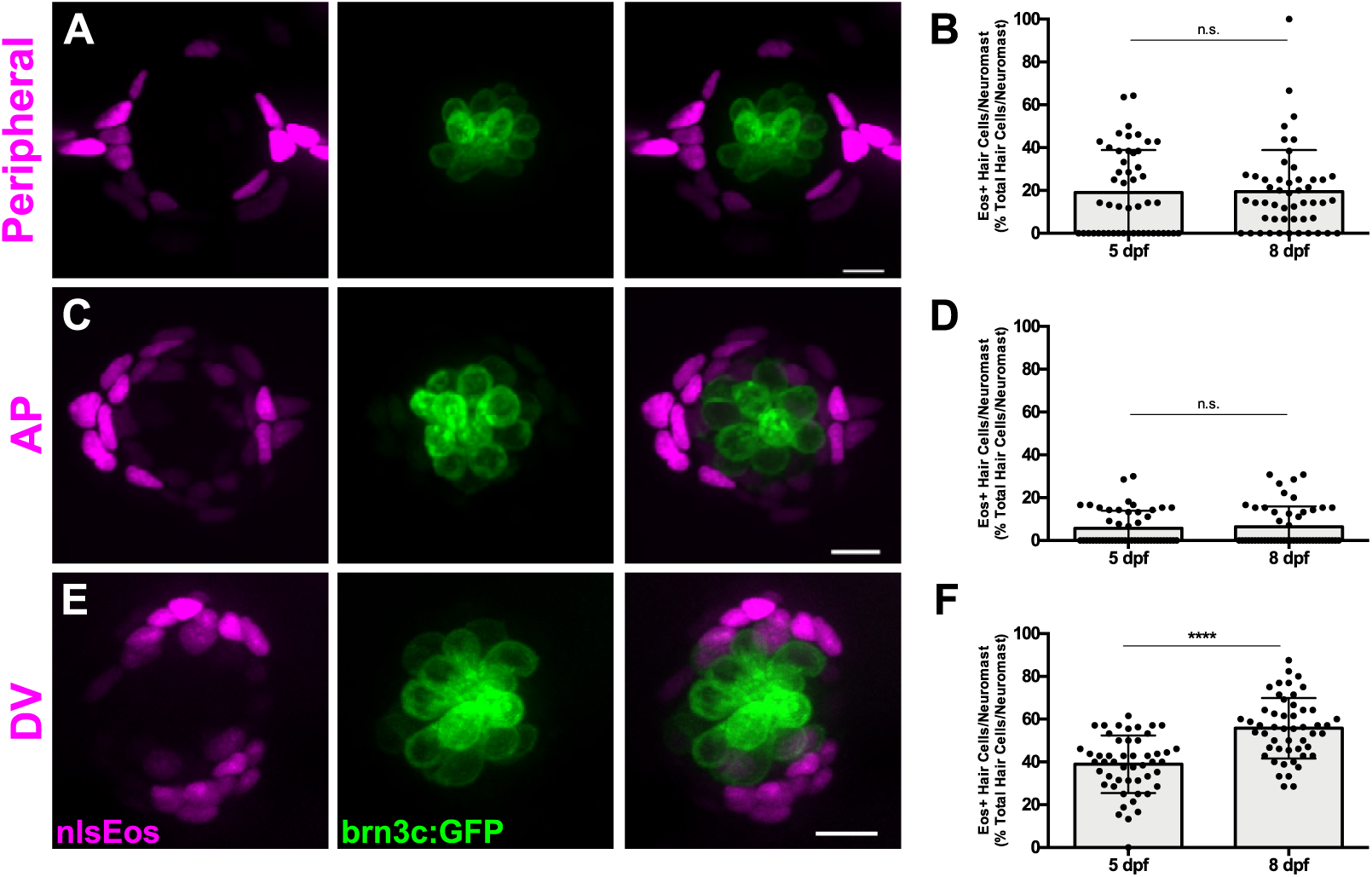
Genetic labeling of spatially-distinct support cell populations. **(A, C, E)** Maximum projections of neuromasts from *sfrp1a*:nlsEos (Peripheral, A), *tnfsf10l3*:nlsEos (AP, C), and *sost*:nlsEos (DV, E) fish. Converted nlsEos-positive cells are shown in magenta, and brn3c:GFP-positive hair cells are shown in green. Scale bar = 10 μm. **(B, D, F)** Percentage of hair cells per neuromast labeled by Peripheral (B), AP (D), and DV cells (F) at 5 and 8 dpf. **(B)** 5 dpf: 19.04 ± 19.86, n = 50 neuromasts; 8 dpf: 19.46 ± 19.44, n = 50 neuromasts; mean ± SD; Mann Whitney U test, p = 0.7047. **(D)** 5 dpf: 5.71 ± 8.22, n = 50 neuromasts; 8 dpf: 6.36 ± 9.57, n = 50 neuromasts; mean ± SD; Mann Whitney U test, p = 0.9668. **(F)** 5 dpf: 38.93 ± 13.46, n = 50 neuromasts; 8 dpf:55.78 ± 14.13, n = 50 neuromasts; mean ± SD; Mann Whitney U test, p < 0.0001.

We first examined how these different support cell populations contributed to hair cell development and turnover under homeostatic conditions. All three nlsEos lines were crossed to a hair cell-specific transgenic line (Tg[Brn3c:GAP43-GFP]^s356t^ (Xiao et al. 2005), hereafter known as brn3c:GFP) in order to distinguish hair cell nuclei. Eos in support cells was photoconverted at 5 dpf and larvae were fixed and immunostained for GFP either immediately or at 8 dpf. At 5 dpf, 19% of hair cells were labeled with Eos expressed by the Peripheral cell transgene, and this number remained the same by 8 dpf (Fig. 2B; p = 0.7047). Eos from the AP cell transgene labeled about 6% of hair cells at both 5 and 8 dpf (Fig. 2D; p = 0.9668). Since there is no change over the three-day span, neither of these populations are responsible for generating new hair cells under homeostatic conditions. In contrast, the amount of hair cells labeled with photoconverted Eos from the DV cell transgene increased from 39% to 56% over that three-day span (Fig. 2F; p < 0.0001). Thus, the DV cell population seems to be predominantly involved in ongoing hair cell generation during homeostasis.

We next used these transgenic lines to determine whether there was any functional difference between these support cell subpopulations regarding their ability to serve as hair cell progenitors during regeneration. Each of the nlsEos lines were once again crossed to brn3c:GFP fish in order to distinguish hair cell nuclei. Eos in support cells was photoconverted at 5 dpf, and larvae were subjected to neomycin-induced hair cell ablation and then fixed and immunostained for GFP at 72 hpt (Fig. 3A). Only 4% of regenerated hair cells were derived from the Peripheral cell population, whereas the AP cell and DV cell populations contributed significantly more, generating 20% and 61% of regenerated hair cells, respectively (Fig. 3B-D, arrowheads; Fig. 3E; p = 0.003 [Peripheral vs. AP], p < 0.0001 [Peripheral/AP vs. DV]). In order to ensure that this difference in Eos incorporation was not simply due to relative proportion of available Eos-positive support cells, we counted the number of Eos-positive support cells in each transgenic line at 5 dpf, prior to hair cell ablation. There were about half as many Peripheral cells relative to the other two populations, but no significant difference between the number of AP cells and the number of DV cells (Fig. 3F; Peripheral = 14.30 ± 4.17; AP = 22.8 ± 4.40; DV = 23.86 ± 4.45; p < 0.0001 [Peripheral vs. AP/DV], p > 0.9999 [AP vs. DV]). Thus, the difference in regenerative capacity between these populations is not simply a reflection of the number of available cells, but rather of differences in the progenitor identity of the populations.

**Figure 3.**
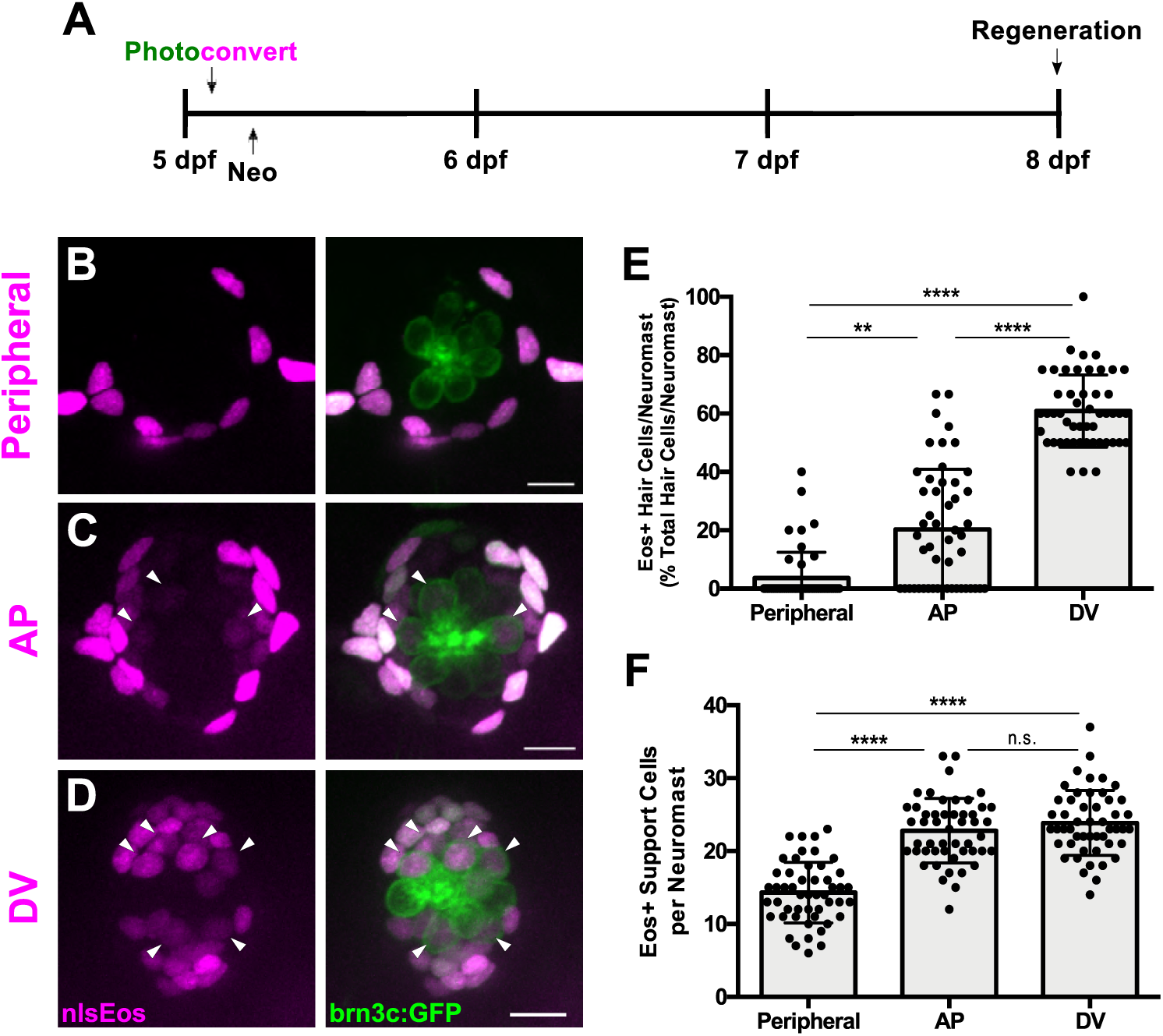
Distinct support cell populations have different regenerative capacities. **(A)** Timeline of nlsEos fate mapping experiment. Fish were photoconverted at 5 dpf, treated with neomycin, then fixed and imaged 72 hours post treatment (8 dpf). **(B, C, D)** Maximum projections of neuromasts from *sfrp1a*:nlsEos (Peripheral, B), *tnfsf10l3*:nlsEos (AP, C), and *sost*:nlsEos (DV, D) fish following photoconversion and hair cell regeneration. Converted nlsEos-positive cells are shown in magenta, and brn3c:GFP-positive hair cells are shown in green. Arrowheads indicate nlsEos-positive hair cells. Scale bar = 10 μm. **(E)** Percentage of hair cells per neuromast labeled by nlsEos following regeneration. *Sfrp1a*:nlsEos (Peripheral): 3.59 ± 8.87, n = 50 neuromasts; *tnfsf10l3*:nlsEos (AP): 20.28 ± 20.58, n = 50 neuromasts; *sost*:nlsEos (DV): 60.87 ± 12.37, n = 50 neuromasts; mean ± SD; Kruskal-Wallis test, Dunn’s post-test, p = 0.003 (Peripheral vs. AP), p < 0.0001 (Peripheral vs. DV, AP vs. DV). **(F)** Total nlsEos-positive support cells per neuromast prior to hair cell ablation. *Sfrp1a*:nlsEos (Peripheral): 14.30 ± 4.17, n = 50 neuromasts; *tnfsf10l3*:nlsEos (AP): 22.8 ±4.40, n = 50 neuromasts; *sost*:nlsEos (DV): 23.86 ± 4.45, n = 50 neuromasts; mean ± SD; Kruskal-Wallis test, Dunn’s post-test, p < 0.0001 (Peripheral vs. AP, Peripheral vs. DV), p > 0.9999 (AP vs. DV).

### Inhibition of Notch Signaling Differentially Impacts Support Cell Subpopulations

Notch-mediated lateral inhibition plays a crucial role in ensuring the proper number of hair cells are regenerated, and inhibition of Notch signaling following hair cell damage dramatically increases the number of regenerated hair cells (Ma, Rubel, and Raible 2008; Wibowo et al. 2011; Romero-Carvajal et al. 2015). Thus, we examined how Notch inhibition impacted the progenitor function of our three support cell populations. We crossed all of our nlsEos lines to the brn3c:GFP line, and treated double-positive larvae with 50 μM LY411575 (LY), a potent γ-secretase inhibitor (Mizutari et al. 2013; Romero-Carvajal et al. 2015), for 24 hours immediately following neomycin treatment. Fish were fixed at 72 hpt and immunostained for GFP. In all three lines, after Notch inhibition (Neo/LY) there were roughly twice as many hair cells as control fish (Neo) (Fig. 4A-B, E-F, I-J; p < 0.0001 [all lines]), consistent with previous studies. The small number of Peripheral cell-derived hair cells was no different between LY-treated fish and non-treated fish (Fig. 4C; 0.62 ± 1.28 [Neo] vs. 1.15 ± 2.16 [Neo/LY]; p = 0.2481). By contrast, the number of nlsEos-positive hair cells from both AP and DV cells increased in fish treated with LY. Moreover, while the number of nlsEos-positive hair cells derived from DV cells doubled in LY-treated fish (Fig. 4K; 7.40 ± 2.13 [Neo] vs. 15.25 ± 6.36 [Neo/LY]; p < 0.0001), those derived from AP cells increased roughly five-fold (Fig. 4G; 2.22 ± 1.94 [Neo] vs. 11.38 ± 4.23 [Neo/LY]; p < 0.0001). As a consequence, the percentage of hair cells derived from DV cells decreased correspondingly (Fig. 4L; 67.86 ± 14.63 [Neo] vs. 54.69 ± 14.01 [Neo/LY]; p < 0.0001), whereas those derived from AP cells doubled (Fig. 4H; 25.19 ± 21.72 [Neo] vs. 50.68 ± 19.23 [Neo/LY]; p < 0.0001). These data suggest that generation of hair cells from both the AP and DV populations is regulated by Notch signaling (with the AP population being regulated to a greater extent), whereas Peripheral cells are not responsive to Notch signaling.

**Figure 4.**
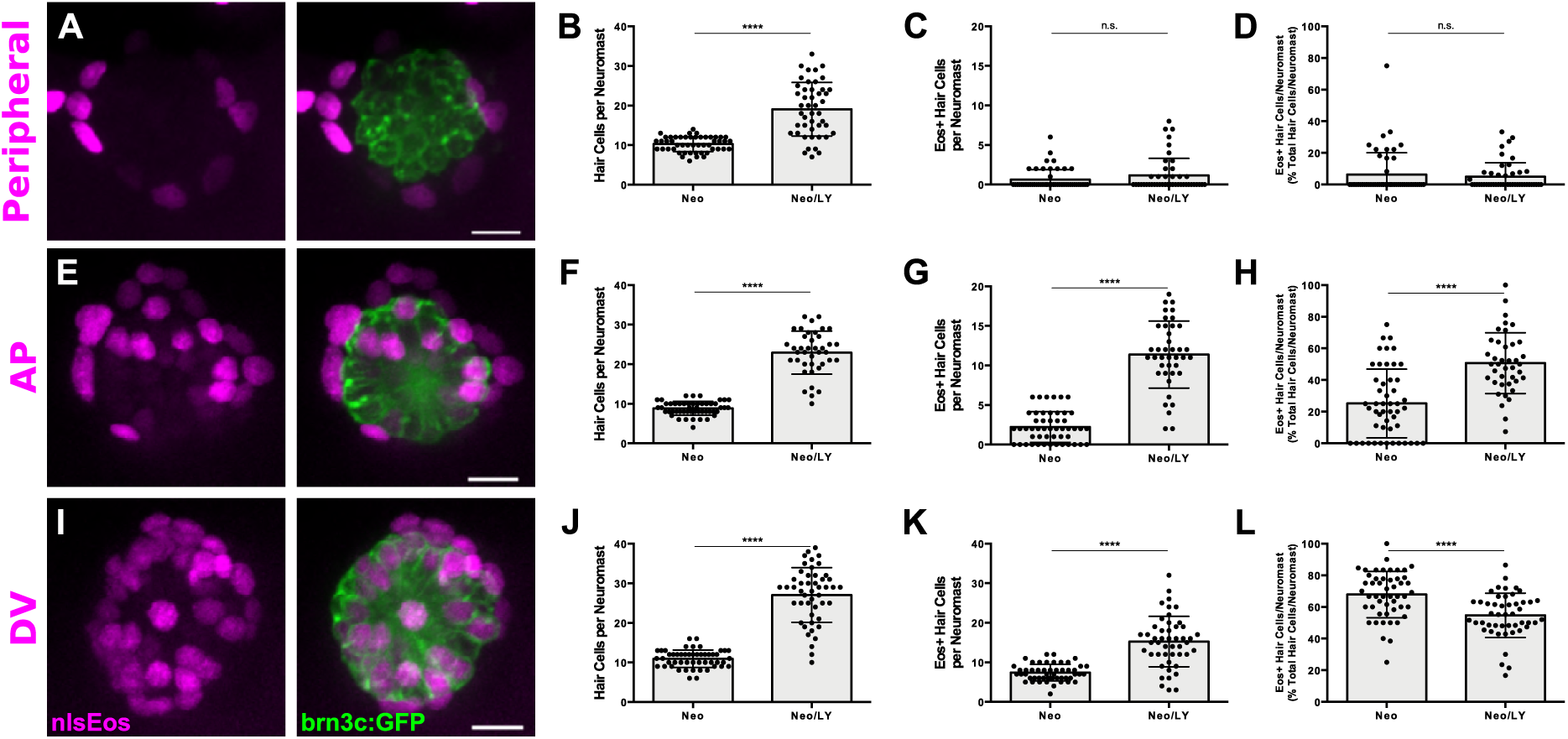
Notch signaling differentially regulates support cell populations. **(A, E, I)** Maximum projections of neuromasts expressing *sfrp1a*:nlsEos (Peripheral, A), *tnfsf10l3*:nlsEos (AP, E), and *sost*:nlsEos (DV, I) following Notch-inhibited hair cell regeneration. Converted nlsEos-positive cells are shown in magenta, and brn3c:GFP-positive hair cells are shown in green. Scale bar = 10 μm. **(B)** Total number of hair cells per neuromast in *sfrp1a*:nlsEos fish following hair cell regeneration. Neo: 10.28 ± 1.88, n = 50 neuromasts; Neo/LY: 19.07 ± 6.79, n = 46 neuromasts; mean ± SD; Mann Whitney U test, p < 0.0001. **(C)** *Sfrp1a*:nlsEos-positive hair cells per neuromast following hair cell regeneration. Neo: 0.62 ± 1.28, n = 50 neuromasts; Neo/LY: 1.15 ± 2.16, n = 46 neuromasts; mean ± SD; Mann Whitney U test, p = 0.2481. **(D)** Percentage of *sfrp1a*:nlsEos-labeled hair cells per neuromast following hair cell regeneration. Neo: 6.31 ± 13.83, n = 50 neuromasts; Neo/LY: 4.95 ± 8.82, n = 46 neuromasts; mean ± SD; Mann Whitney U test, p = 0.5148. **(F)** Total number of hair cells per neuromast in *tnfsf10l3*:nlsEos fish following hair cell regeneration. Neo: 8.84 ± 1.75, n = 50 neuromasts; Neo/LY: 22.93 ± 5.45, n = 40 neuromasts; mean ± SD; Mann Whitney U test, p < 0.0001. **(G)** *Tnfsf10l3*:nlsEos-positive hair cells per neuromast following hair cell regeneration. Neo: 2.22 ± 1.94, n = 50 neuromasts; Neo/LY: 11.38 ± 4.23, n = 40 neuromasts; mean ± SD; Mann Whitney U test, p < 0.0001. **(H)** Percentage of *tnfsf10l3*:nlsEos-labeled hair cells per neuromast following hair cell regeneration. Neo: 25.19 ± 21.72, n = 50 neuromasts; Neo/LY: 50.68 ± 19.23, n = 40 neuromasts; mean ± SD; Mann Whitney U test, p < 0.0001. **(J)** Total number of hair cells per neuromast in *sost*:nlsEos fish following hair cell regeneration. Neo: 10.94 ± 2.23, n = 50 neuromasts; Neo/LY: 27.06 ± 6.90, n = 48 neuromasts; mean ± SD; Mann Whitney U test, p < 0.0001. **(K)** *Sost*:nlsEos-positive hair cells per neuromast following hair cell regeneration. Neo: 7.40 ± 2.13, n = 50 neuromasts; Neo/LY: 15.25 ± 6.36, n = 48 neuromasts; mean ± SD; Mann Whitney U test, p < 0.0001. **(L)** Percentage of *sost*:nlsEos-labeled hair cells per neuromast following hair cell regeneration. Neo: 67.86 ± 14.63, n = 50 neuromasts; Neo/LY: 54.69 ± 14.01, n = 48 neuromasts; mean ± SD; Mann Whitney U test, p < 0.0001.

### Selective Ablation of DV Cells Reduces Hair Cell Regeneration

Since the DV cell population generates roughly 60% of regenerated hair cells, we sought to determine whether these cells were required for hair cell regeneration. To this end, we generated a transgenic line in which an enhanced-potency nitroreductase (epNTR; Tabor et al., 2014) fused to GFP was introduced into the *sost* locus using CRISPR (Tg[*sost*:epNTR-GFP]^w216^, hereafter known as *sost*:NTR-GFP). Nitroreductase is a bacterial enzyme that selectively binds its prodrug Metronidazole (Mtz), converting Mtz into toxic metabolites that kill the cells expressing it (Curado et al. 2007). We then compared the extent of *sost:NTR-GFP* expression in DV cells, as defined by the *sost:nlsEos* transgene. At 3 dpf, soon after the initiation of transgene expression, we see considerable overlap between NTR-GFP and nlsEos. All NTR-GFP+ cells were also positive for nlsEos, while an additional subset of cells expressed nlsEos alone. When we compared expression at 5 dpf, the size of the double-positive (NTR-GFP+; nlsEos+) population did not change, whereas the number of cells expressing nlsEos alone increased significantly, occupying a more central location (Fig. 5A-B, arrowheads; Fig. 5C; NTR-GFP/nlsEos: 9.04 ± 2.39 [3 dpf] vs. 8.47 ± 2.27 [5 dpf]; nlsEos only: 6.10 ± 2.27 [3 dpf] vs.10.86 ± 2.72 (5 dpf); p > 0.9999 [NTR-GFP/nlsEos], p < 0.0001 [nlsEos only]). These observations are consistent with the idea that both transgenes initiate expression at the same time, but that nlsEos protein is retained longer than NTR-GFP protein as cells mature and as a result, NTR-GFP is expressed in a subset of DV cells. We next tested to the efficacy of DV cell ablation at 3 and 5 dpf. Treatment of these fish with 10 mM Mtz for 8 hours was sufficient to ablate the majority of NTR-GFP cells. Treating fish with Mtz for 8 hours at 5 dpf (Mtz5) slightly but significantly decreased the number of support cells solely expressing nlsEos by about 13%. Treating fish with Mtz for 8 hours at 3 dpf, followed by a second 8-hour Mtz treatment at 5 dpf (Mtz3/5) decreased the number of solely nlsEos-positive cells even further, by about 40% (Fig. 5D-G; Mock: 11.18 ± 2.04; Mtz5: 9.72 ± 2.03; Mtz3/5: 6.76 ± 2.12; p = 0.0288 [Mock vs. Mtz5], p < 0.0001 [Mock vs. Mtz3/5, Mtz5 vs. Mtz3/5]).

**Figure 5.**
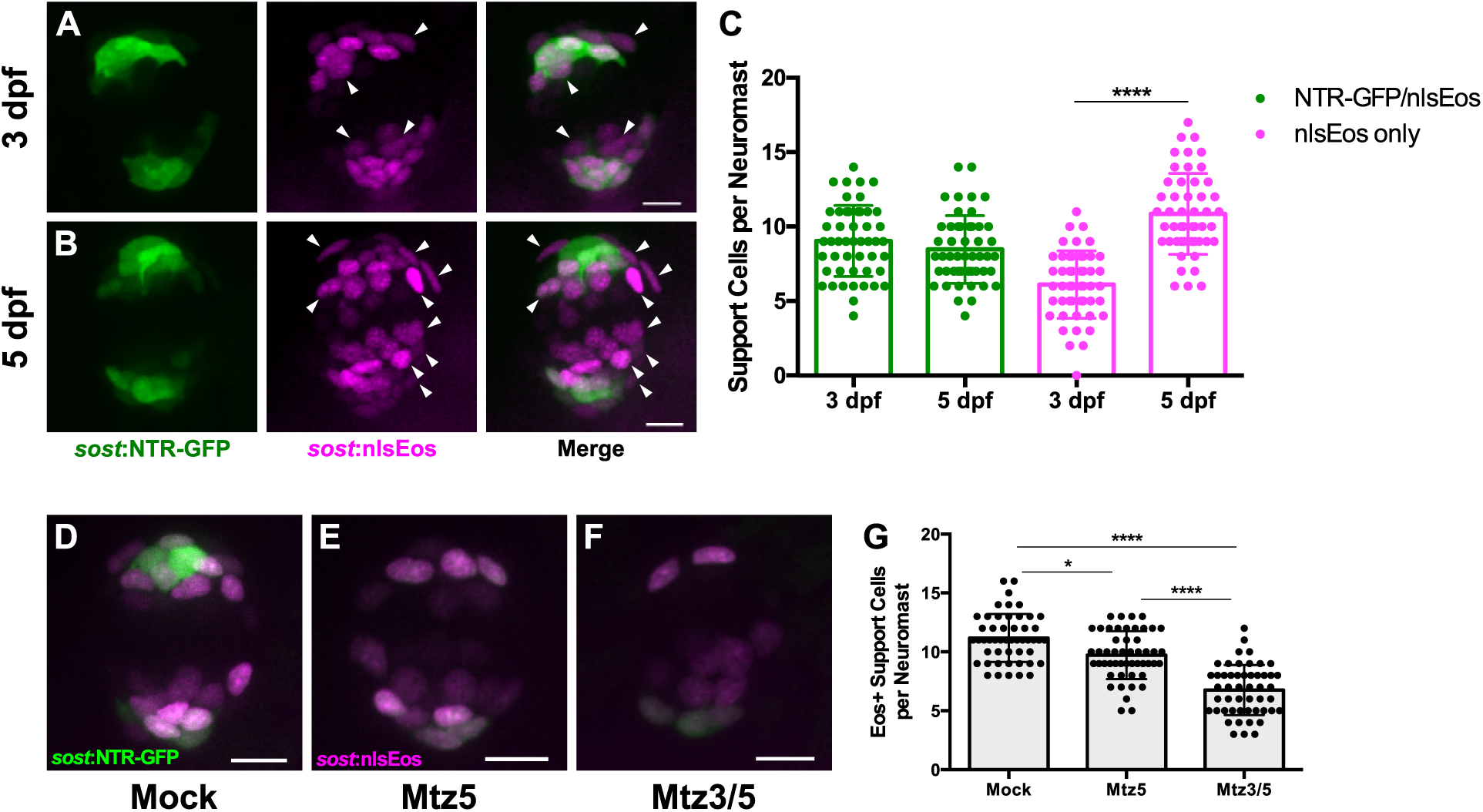
Differences in overlap between *sost*:NTR-GFP and *sost*:nlsEos populations. **(A-B)** Maximum projections of neuromasts from *sost*:NTR-GFP; *sost*:nlsEos fish at 3 dpf (A) and 5 dpf (B). *Sost*:NTR-GFP cells are shown in green and *sost*:nlsEos cells are shown in magenta. Arrowheads indicate cells expressing *sost*:nlsEos but not *sost*:NTR-GFP. Scale bar = 10 μm. **(C)** Support cells per neuromast expressing either NTR-GFP and nlsEos (green) or nlsEos only (magenta) at 3 dpf and 5 dpf. NTR-GFP/nlsEos: 9.04 ± 2.39 (3 dpf) vs. 8.47 ± 2.27 (5 dpf), n = 49 neuromasts; nlsEos only: 6.10 ± 2.27 (3 dpf) vs. 10.86 ± 2.72 (5 dpf), n = 49 neuromasts; mean ± SD; Kruskal-Wallis test, Dunn’s post-test, p > 0.9999 (NTR-GFP/nlsEos 3 dpf vs. 5 dpf), p < 0.0001 (nlsEos only 3 dpf vs. 5 dpf). **(D-F)** Maximum projections of neuromasts from *sost*:NTR-GFP; *sost*:nlsEos fish following mock treatment (D; Mock), Mtz at 5 dpf (E; Mtz5), and Mtz at 3 dpf and 5 dpf (F; Mtz3/5). *Sost*:NTR-GFP cells are shown in green and *sost*:nlsEos cells are shown in magenta. Scale bar = 10 μm. **(G)** Support cells per neuromast solely expressing *sost*:nlsEos following Mtz treatment. Mock: 11.18 ± 2.04, n = 50 neuromasts; Mtz5: 9.72 ± 2.03, n = 50 neuromasts; Mtz3/5: 6.76 ± 2.12, n = 50 neuromasts; mean ± SD; Kruskal-Wallis test, Dunn’s post-test, p = 0.0288 (Mock vs. Mtz5), p < 0.0001 (Mock vs. Mtz3/5, Mtz5 vs. Mtz3/5).

We next tested the impact of DV cell ablation on hair cell regeneration. We compared two groups: neomycin exposure followed by Mtz treatment at 5 dpf (Neo/Mtz5), compared to Mtz treatment at 3 dpf, then neomycin treatment at 5 dpf followed by a second Mtz treatment (Mtz3/Neo/Mtz5; Fig. 6A). For both groups, nlsEos was photoconverted at 5 dpf, just prior to neomycin treatment, and larvae were fixed at 72 hpt and immunostained for GFP and Parvalbumin (to label NTR-GFP+ cells and hair cells, respectively). The Neo/Mtz5 treatment resulted in a small but significant reduction in both hair cells and nlsEos-positive hair cells per neuromast relative to normal regeneration (Fig. 6B-C, E-F; Total hair cells: 11.73 ± 2.10 [Neo] vs. 9.33 ± 1.88 [Neo/Mtz5]; p = 0.0001; nlsEos+ hair cells: 7.78 ± 2.36 [Neo] vs. 4.90 ± 2.02 [Neo/Mtz5]; p = 0.0003). The Mtz3/Neo/Mtz5-treated larvae exhibited even fewer hair cells per neuromast (Fig. 6D, E; 11.73 ± 2.10 [Neo] vs. 7.52 ± 1.74 [Mtz3/Neo/Mtz5]; p < 0.0001), with nlsEos-labeled hair cells decreased to a mere 14% of total regenerated hair cells (Fig. 6G; 65.81 ± 14.89 [Neo] vs. 14.29 ± 18.10 [Mtz3/Neo/Mtz5]; p < 0.0001). Importantly, Mtz treatment of siblings without the *sost:*NTR-GFP transgene had no impact on hair cell regeneration (Fig. 6 – figure supplement 1; Neo: 9.5 ± 1.50; Mtz3/Neo/Mtz5: 9.98 ± 1.51; p = 0.2317). Thus, ablation of DV cells reduces the number of hair cells regenerated.

**Figure 6.**
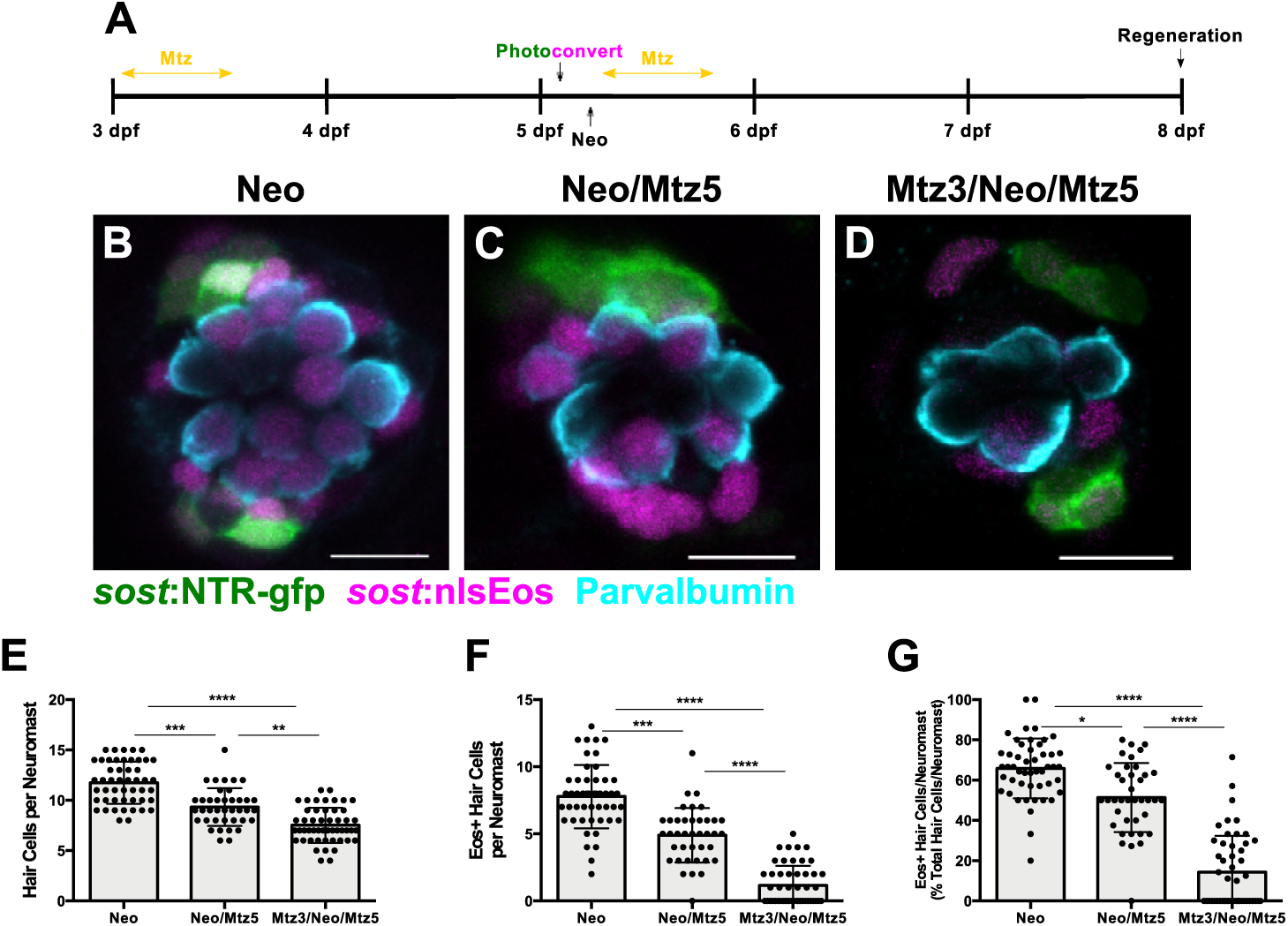
Ablation of DV cells decreases number of regenerated hair cells. **(A)** Timeline of DV cell-ablation experiment. Larvae were treated with Mtz at 3 dpf, photoconverted, then treated with neomycin, then treated with Mtz again at 5 dpf, and fixed and immunostained at 72 hpt (8 dpf). **(B-D)** Maximum projections of neuromasts from *sost*:NTR-GFP; *sost*:nlsEos fish following neomycin (B; Neo), neomycin and Mtz (C; Neo/Mtz5), and Mtz, neomycin, and Mtz treatments (D; Mtz3/Neo/Mtz5). *Sost*:NTR-GFP cells are shown in green, *sost*:nlsEos cells are shown in magenta, and anti-Parvalbumin-stained hair cells are shown in cyan. Scale bar = 10 μm. **(E)** Total hair cells per neuromast following regeneration. Neo: 11.73 ± 2.10, n = 49 neuromasts; Neo/Mtz5: 9.33 ± 1.88, n = 39 neuromasts; Mtz3/Neo/Mtz5: 7.52 ± 1.74, n = 50 neuromasts; mean ± SD; Kruskal-Wallis test, Dunn’s post-test, p = 0.0001 (Neo vs. Neo/Mtz5), p < 0.0001 (Neo vs. Mtz3/Neo/Mtz5), p = 0.0016 (Neo/Mtz5 vs. Mtz3/Neo/Mtz5). **(F)** *Sost*:nlsEos-positive hair cells per neuromast following regeneration. Neo: 7.78 ± 2.36, n = 49 neuromasts; Neo/Mtz5: 4.90 ± 2.02, n = 39 neuromasts; Mtz3/Neo/Mtz5: 1.16 ± 1.46, n = 50 neuromasts; mean ± SD; Kruskal-Wallis test, Dunn’s post-test, p = 0.0003 (Neo vs. Neo/Mtz5), p < 0.0001 (Neo vs. Mtz3/Neo/Mtz5, Neo/Mtz5 vs. Mtz3/Neo/Mtz5). **(G)** Percentage of hair cells per neuromast labeled by *sost*:nlsEos following regeneration. Neo: 65.81 ± 14.89, n = 49 neuromasts; Neo/Mtz5: 51.40 ± 17.17, n = 39 neuromasts; Mtz3/Neo/Mtz5: 14.29 ± 18.10, n = 50 neuromasts; mean ± SD; Kruskal-Wallis test, Dunn’s post-test, p = 0.0147 (Neo vs. Neo/Mtz5), p < 0.0001 (Neo vs. Mtz3/Neo/Mtz5, Neo/Mtz5 vs. Mtz3/Neo/Mtz5).

We then examined how Notch signaling impacted hair cell regeneration in the context of DV cell ablation. We treated *sost*:NTR-GFP larvae with 50 μM LY for 24 hours following ablation (Mtz3/Neo/Mtz5/LY) and assayed hair cell number at 72 hpt (Fig. 7A). As expected, the number of regenerated hair cells increased significantly after LY treatment in all groups (Fig.7B-F; p < 0.0001), and DV cell ablation significantly decreased hair cell regeneration (Fig. 7B, D, F; 9.42 ± 1.85 [Neo] vs. 6.86 ± 1.76 [Mtz3/Neo/Mtz5]; p = 0.0058). However, LY treatment following Mtz ablation resulted in significantly fewer regenerated hair cells than LY alone (Fig. 7C, E, F; 21.08 ± 4.42 [Neo/LY] vs. 15.06 ± 3.51 [Mtz3/Neo/Mtz5/LY]; p = 0.0029), indicating that inhibiting Notch signaling cannot fully compensate for the loss of the DV population.

**Figure 7.**
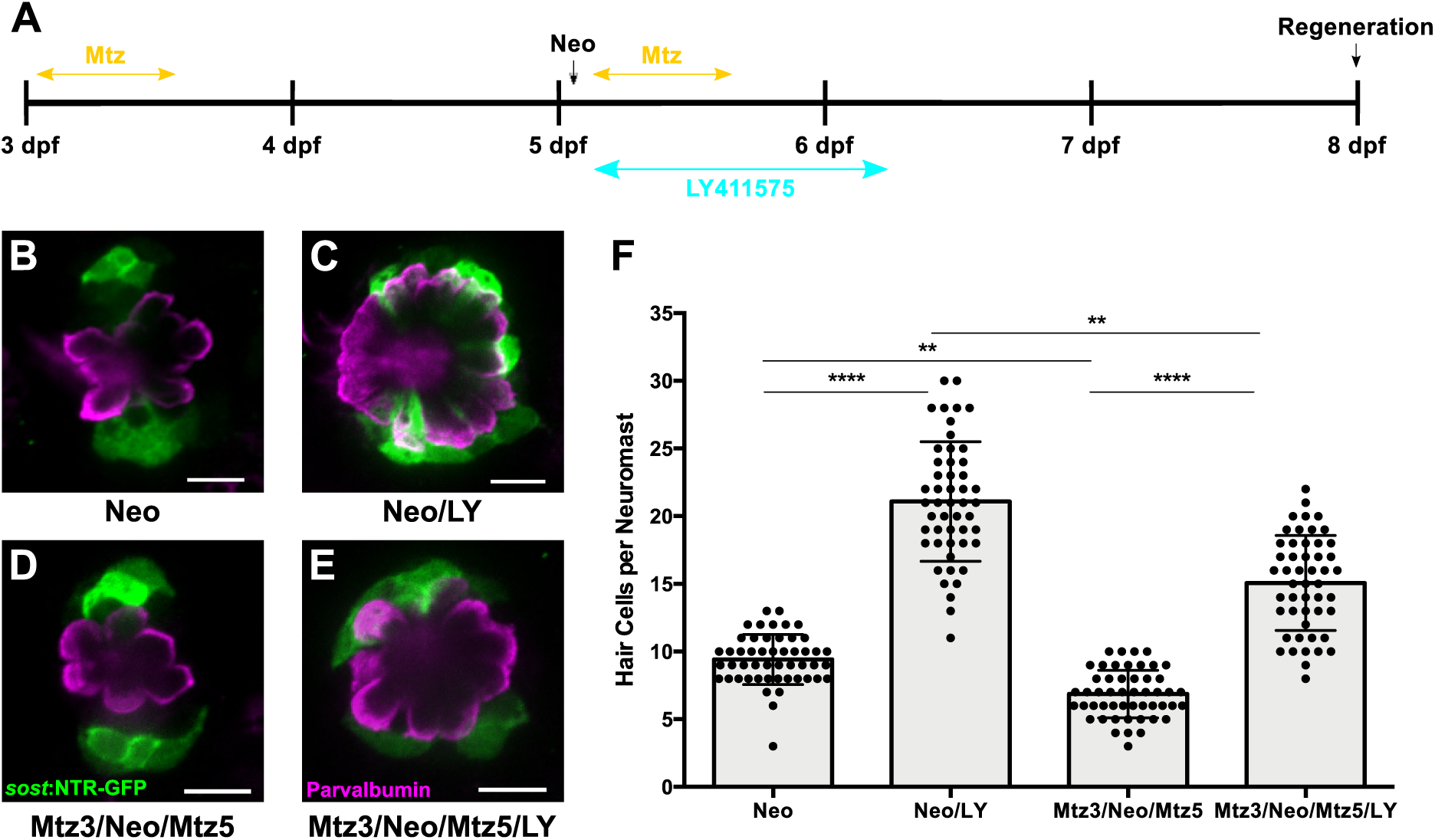
DV cell-ablation reduces the number of supernumerary hair cells formed during Notch-inhibited hair cell regeneration. **(A)** Timeline of dual DV cell-ablation, Notch-inhibition experiment. *Sost*:NTR-GFP larvae were treated with Mtz at 3 dpf, treated with neomycin at 5dpf, then co-treated with Mtz and LY411575 for 8 hours, then washed out and treated with LY411575 for 16 additional hours (24 hours total LY). **(B-E)** Maximum projections of *sost*:NTR-GFP neuromasts following normal hair cell regeneration (B; Neo), Notch-inhibited hair cell regeneration (C; Neo/LY), DV cell-ablated hair cell regeneration (D; Mtz3/Neo/Mtz5), and DV cell-ablated and Notch-inhibited hair cell regeneration (E; Mtz3/Neo/Mtz5/LY). *Sost*:NTR-GFP cells are shown in green, and anti-Parvalbumin immunostained hair cells are shown in magenta. Scale bar = 10 μm. **(F)** Total number of hair cells per neuromast following hair cell regeneration. Neo: 9.42 ± 1.85, n = 50 neuromasts; Neo/LY: 21.08 ± 4.42, n = 50 neuromasts; Mtz3/Neo/Mtz5: 6.86 ± 1.76, n = 50 neuromasts; Mtz3/Neo/Mtz5/LY: 15.06 ± 3.51, n = 50 neuromasts; mean ± SD; Kruskal-Wallis test, Dunn’s post-test, p < 0.0001 (Neo vs. Neo/LY; Mtz3/Neo/Mtz5 vs. Mtz3/Neo/Mtz5/LY), p = 0.0058 (Neo vs. Mtz3/Neo/Mtz5), p = 0.0029 (Neo/LY vs. Mtz3/Neo/Mtz5/LY).

### AP and DV Cells Define Separate Progenitor Populations

While the DV population generates roughly 60% of hair cells after damage, the other 40% must derive from a different population. Consistent with this observation, reduction of DV cells by Mtz treatment only partially blocks new hair cell formation, indicating that there must be additional progenitor populations. We believed that AP cells could define this additional population, since they were capable of generating roughly 20% of regenerated hair cells (Fig.3E). However, there may be some overlap between the expression of *tnfsf10l3*:nlsEos defining the AP domain and *sost*:nlsEos defining the DV domain. When we crossed the *tnfsf10l3*:nlsEos and *sost*:nlsEos lines together, we found that roughly 88% of regenerated hair cells were nlsEos positive when larvae expressed both transgenes, compared to 65% from *sost*:nlsEos alone and 28% from *tnfsf10l3*:nlsEos alone (Fig. 8A-E; p < 0.0001). Thus, while not completely additive, these data suggest that the AP population is distinct from the DV population in terms of its progenitor function.

**Figure 8.**
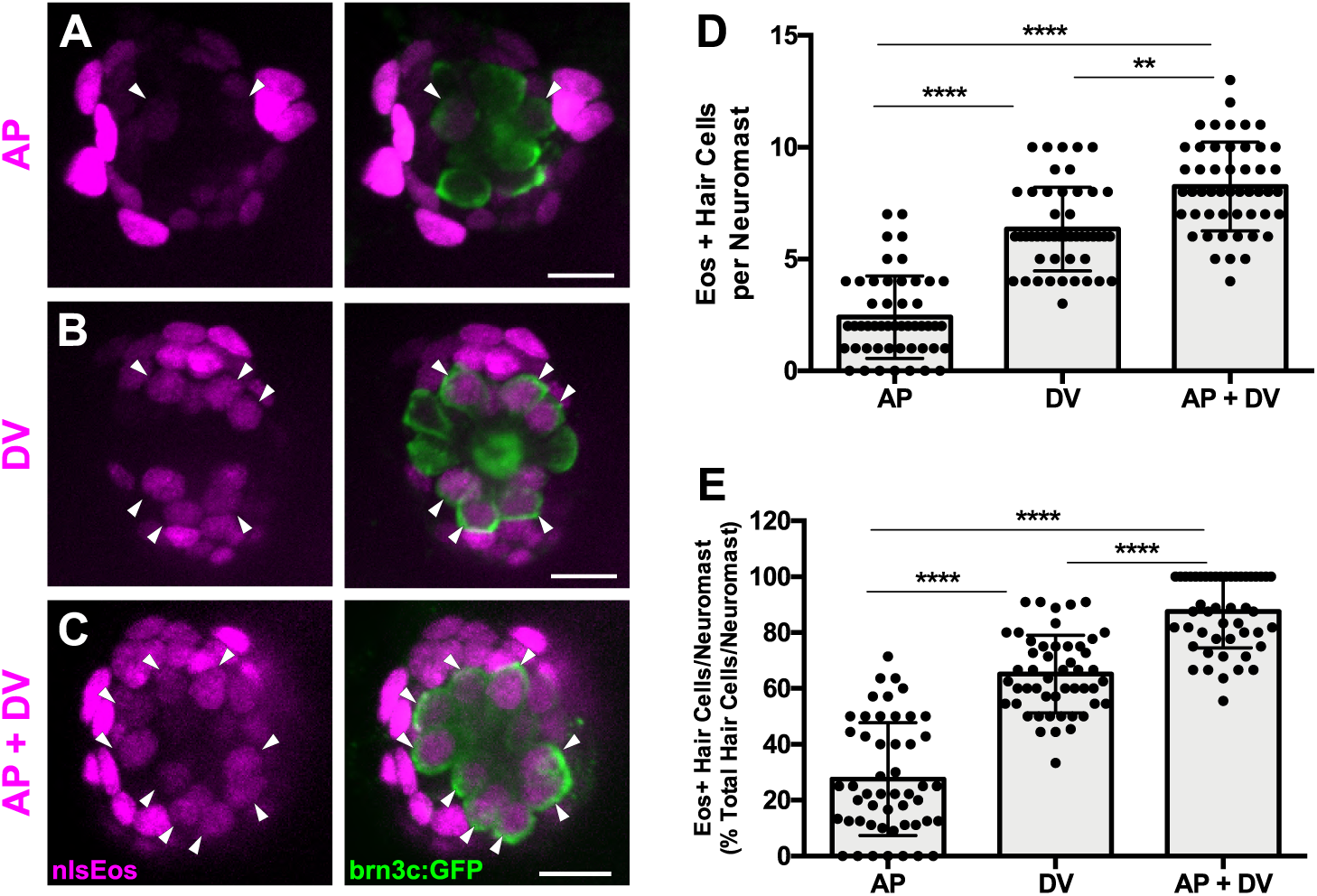
AP cells and DV cells define separate progenitor populations. **(A-C)** Maximum projections of neuromasts from *tnfsf10l3*:nlsEos (AP, A), *sost*:nlsEos (DV, B), and *tnfsf10l3*:nlsEos/*sost*:nlsEos fish (AP + DV, C) following photoconversion and regeneration. Converted nlsEos-positive cells are shown in magenta, and brn3c:GFP-positive hair cells are shown in green. Arrowheads indicate nlsEos-positive hair cells. Scale bar = 10 μm. **(D)** Number of nlsEos-positive hair cells per neuromast in each of the nlsEos lines following regeneration. *Tnfsf10l3*:nlsEos (AP): 2.4 ± 1.84, n = 50 neuromasts; *sost*:nlsEos (DV): 6.34 ± 1.87, n = 50 neuromasts; *tnfsf10l3*:nlsEos/*sost*:nlsEos (AP + DV): 8.24 ± 1.99, n = 50 neuromasts; mean ± SD; Kruskal-Wallis test, Dunn’s post-test, p < 0.0001 (AP vs. DV, AP vs. AP + DV), p = 0.0031 (DV vs. AP + DV). **(E)** Percentage of hair cells per neuromast labeled by nlsEos lines following regeneration. AP: 27.59 ± 20.21, n = 50 neuromasts; DV: 65.16 ± 13.89, n = 50 neuromasts; AP + DV: 87.57 ± 13.02, n = 50 neuromasts; mean ± SD; Kruskal-Wallis test, Dunn’s post-test, p < 0.0001 (all comparisons).

We next examined how the AP population would respond to the ablation of the DV population. We crossed the *tnfsf10l3*:nlsEos line to the *sost*:NTR-GFP line, sorted out double-positive larvae, and compared normal regeneration to that after Mtz treatment (Mtz3/Neo/Mtz5, since this had served to be the best treatment paradigm). As above, nlsEos was photoconverted at 5 dpf, immediately prior to neomycin treatment, and larvae were fixed at 72 hpt and immunostained for GFP and Parvalbumin. Mtz-ablated larvae had significantly fewer hair cells than non-ablated larvae, as in previous experiments (Fig. 9C; 10.36 ± 1.60 [Neo] vs. 7.98 ± 1.74 [Mtz3/Neo/Mtz5]; p < 0.0001), but the number of nlsEos-postive hair cells was no different between the two groups (Fig. 9A-B, arrowheads; Fig. 9D; 2.88 ± 1.83 [Neo] vs. 3.14 ± 1.43 [Mtz3/Neo/Mtz5]; p = 0.3855). The percentage of Eos-positive hair cells did increase, but this is only because the total number of hair cells decreased overall (Fig. 9E; 27.26 ± 16.00 [Neo] vs. 40.43 ± 19.44 [Mtz3/Neo/Mtz5]; p = 0.0002). Thus, the AP population’s progenitor function remains unchanged following DV ablation, providing further support that it is a separate progenitor population from the DV population.

**Figure 9.**
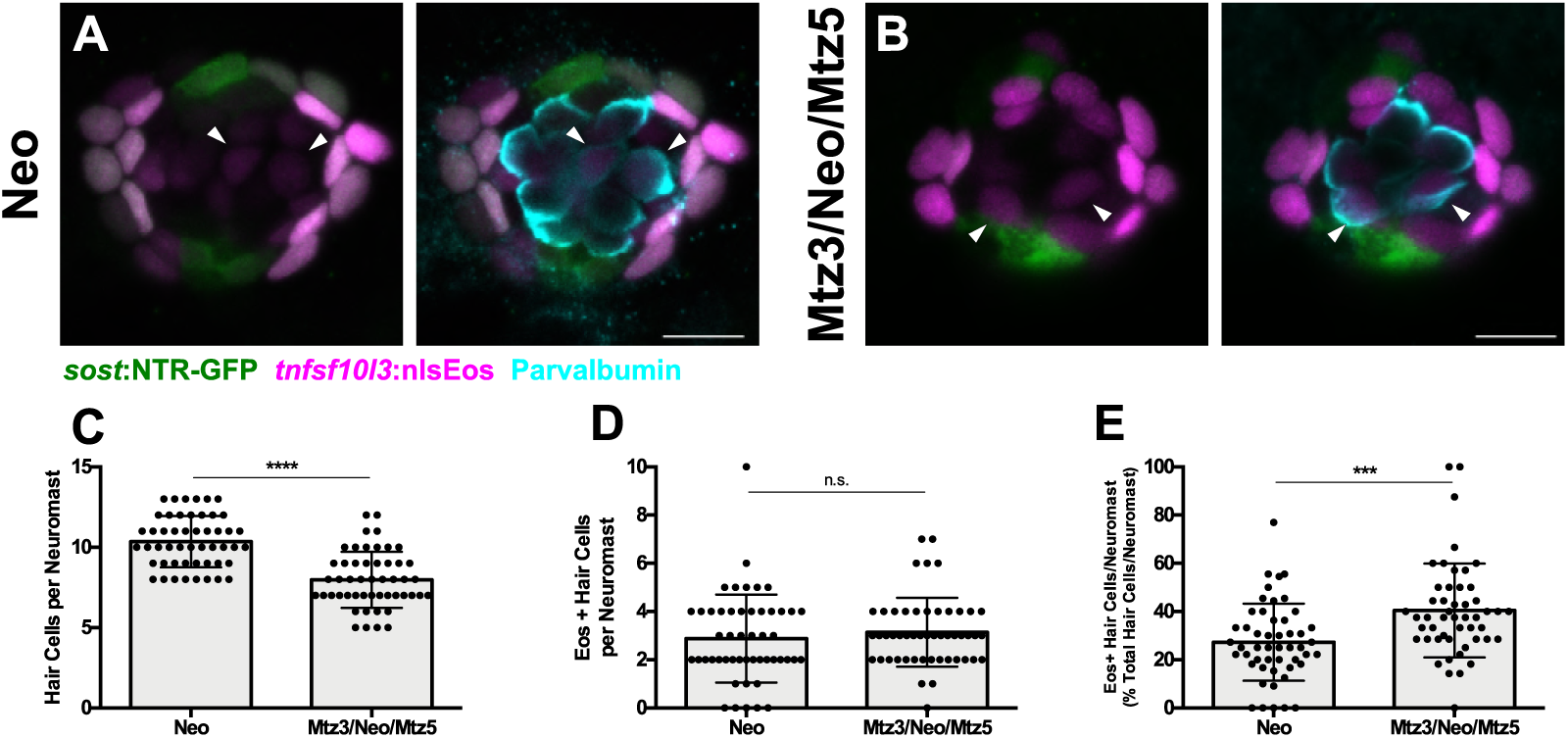
AP population doesn’t compensate for the loss of the DV population during hair cell regeneration. **(A-B)** Maximum projections of *tnfsf10l3*:nlsEos; *sost*:NTR-GFP neuromasts following normal hair cell regeneration (A; Neo) or DV cell-ablated hair cell regeneration (B; Mtz3/Neo/Mtz5). *Sost*:NTR-GFP cells are shown in green, *tnfsf10l3*:nlsEos cells are shown in magenta, and anti-Parvalbumin-stained hair cells are shown in cyan. Arrowheads indicate nlsEos-positive hair cells. Scale bar = 10 μm. **(C)** Total number of hair cells per neuromast following hair cell regeneration. Neo: 10.36 ± 1.60, n = 50 neuromasts; Mtz3/Neo/Mtz5: 7.98 ± 1.74, n = 50 neuromasts; mean ± SD; Mann Whitney U test, p < 0.0001. **(D)** Number of nlsEos-positive hair cells per neuromast following hair cell regeneration. Neo: 2.88 ± 1.83, n = 50 neuromasts; Mtz3/Neo/Mtz5: 3.14 ± 1.43, n = 50 neuromasts; mean ± SD; Mann Whitney U test, p = 0.3855. **(E)** Percentage of hair cells per neuromast labeled by nlsEos following hair cell regeneration. Neo: 27.26 ± 16.00, n = 50 neuromasts; Mtz3/Neo/Mtz5: 40.43 ± 19.44, n = 50 neuromasts; mean ± SD; Mann Whitney U test, p = 0.0002.

### The DV Population Regenerates from Other Support Cell Subpopulations

When examining hair cell regeneration following DV cell ablation, we consistently noticed that there was an increase in NTR-GFP+ cells at 72 hpt. This led us to hypothesize that DV cells were capable of regeneration even in the absence of hair cell damage. To test this, we first administered a 48-hour pulse of EdU (changing into fresh EdU solution after the first 24 hours) immediately following Mtz ablation at 5 dpf and fixed immediately after EdU washout. At 48 hours post ablation, we observed slightly more than half the number of the NTR-GFP+ cells relative to unablated larvae (Fig. 10C; 8.94 ± 1.62 [Mock] vs. 5.34 ± 2.14 [Mtz]; p < 0.0001). However, 58% of NTR-GFP+ cells were EdU-positive in fish treated with Mtz, compared to just 15% in unablated larvae (Fig. 10A-B, arrowheads; Fig. 10D; p < 0.0001). These results indicate that new DV cells arise from proliferation.

**Figure 10.**
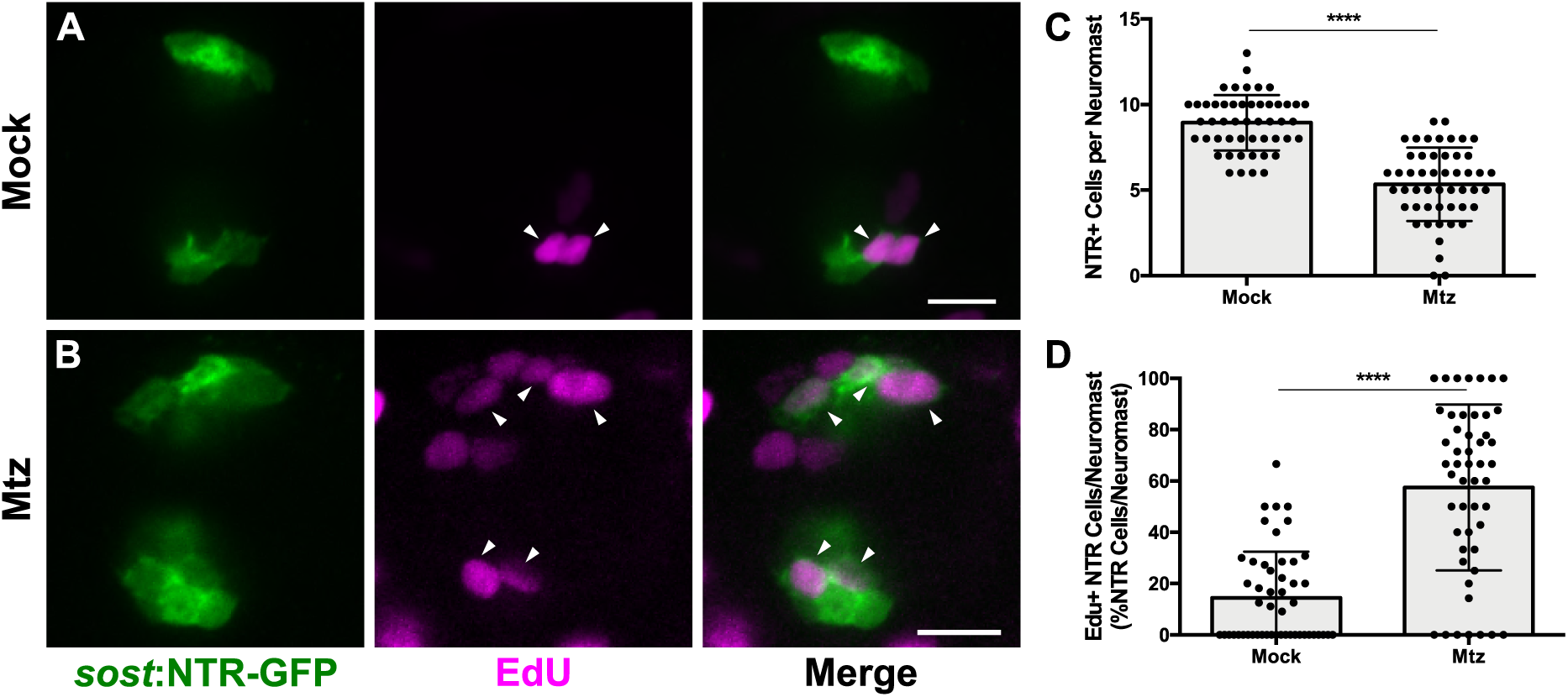
DV population regenerates via proliferation. **(A-B)** Maximum projections of neuromasts from *sost*:NTR-GFP fish either untreated (A; Mock) or treated with 10 mM Mtz (B; Mtz). *Sost*:NTR-GFP cells are shown in green and EdU-positive cells are shown in magenta. Arrowheads indicate EdU-positive *sost*:NTR-GFP cells. Scale bar = 10 μm. **(C)** Total number of *sost*:NTR-GFP cells per neuromast following DV cell regeneration. Mock: 8.94 ± 1.62, n = 50 neuromasts; Mtz: 5.34 ± 2.14, n = 50 neuromasts; mean ± SD; Mann Whitney U test, p < 0.0001. **(D)** Percentage of *sost*:NTR-GFP cells per neuromast labeled by EdU following DV cell regeneration. Mock: 14.47 ± 17.95, n = 50 neuromasts; Mtz: 57.49 ± 32.34, n = 50 neuromasts; mean ± SD; Mann Whitney U test, p < 0.0001.

To determine the source of new DV cells, we crossed *sost*:NTR-GFP fish to our three different nlsEos lines. Double-transgenic fish were photoconverted at 5 dpf, Mtz-ablated, and then fixed at 72 hpt and immunostained for GFP. Following ablation, 56% of NTR-GFP+ cells expressed photoconverted nlsEos when DV cells were labeled, compared to 97% in unablated controls (Fig. 11A-B, arrowheads; Fig. 11C; p < 0.0001). 31% of NTR-GFP+ cells expressed photoconverted nlsEos when Peripheral cells were labeled, compared to 6% in controls (Fig.11D-E, arrowheads; Fig. 11F; p < 0.0001) and 21% of NTR-GFP+ cells expressed photoconverted nlsEos when AP cells were labeled, compared to 7% in controls (Fig. 11G-H, arrowheads; Fig. 11I; p = 0.0004). Thus, DV cells are capable of being replenished after Mtz ablation by other DV cells as well as by both AP and Peripheral cells.

**Figure 11.**
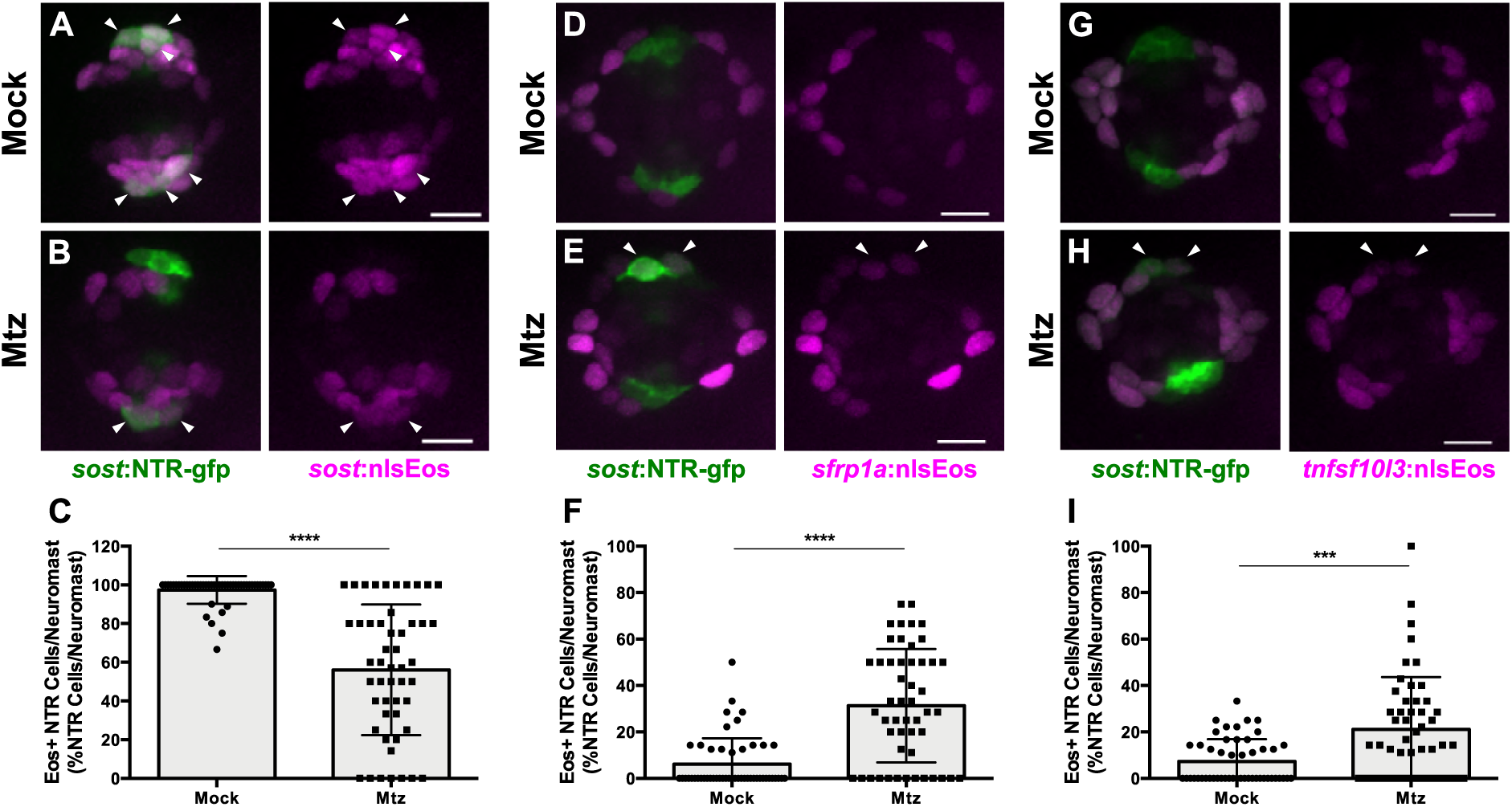
DV cells are replenished by other support cell populations. **(A-B, D-E, G-H)** Maximum projections of neuromasts expressing *sost*:NTR-GFP and *sost*:nlsEos (A-B), *sfrp1a*:nlsEos (D-E), and *tnfsf10l3*:nlsEos (G-H) in the absence of (A, D, G; Mock) or following Mtz-induced DV cell ablation (B, E, H; Mtz). *Sost*:NTR-GFP cells are shown in green and nlsEos-positive cells are shown in magenta. Arrowheads indicate nlsEos-positive *sost*:NTR-GFP cells. Scale bar = 10 μm. **(C)** Percentage of *sost*:NTR-GFP cells per neuromast labeled by *sost*:nlsEos following DV cell regeneration. Mock: 97.39 ± 7.14, n = 50 neuromasts; Mtz: 56.09 ± 33.72, n = 50 neuromasts; mean ± SD; Mann Whitney U test, p < 0.0001. **(F)** Percentage of *sost*:NTR-GFP cells per neuromast labeled by *sfrp1a*:nlsEos following DV cell regeneration. Mock: 6.15 ± 11.14, n = 50 neuromasts; Mtz: 31.27 ± 24.41, n = 50 neuromasts; mean ± SD; Mann Whitney U test, p < 0.0001.**(I)** Percentage of *sost*:NTR-GFP cells per neuromast labeled by *tnfsf10l3*:nlsEos following DV cell regeneration. Mock: 7.31 ± 9.55, n = 50 neuromasts; Mtz: 21.11 ± 22.51, n = 50 neuromasts; mean ± SD; Mann Whitney U test, p = 0.0004.

## DISCUSSION

### Differences in Hair Cell Progenitor Identity Among Support Cell Populations

The data shown above indicate that there are at least three spatially and functionally distinct progenitor populations within the neuromast: (1) a highly regenerative, dorsoventral (DV) population, marked by *sost*:nlsEos, which generates the majority of regenerated hair cells; (2) an anteroposterior (AP) population, marked by *tnfsf10l3*:nlsEos, which also contributes to hair cell regeneration albeit to a far lesser extent than *sost*; and (3) a peripheral population (Peripheral), marked by *sfrp1a*:nlsEos, that does not contain hair cell progenitors (Fig. 12). This model of high regenerative capacity in the dorsoventral region and low regenerative capacity in the anteroposterior region is consistent with the label-retaining studies performed by Cruz et al., (2015) as well as the BrdU-localization studies of Romero-Carvajal et al., (2015). However, an examination of the overlap in expression between *sost*:nlsEos and *sost*:NTR-GFP reveals distinctions even amongst this DV progenitor population. We hypothesize that cells that express only nlsEos have matured from those that express both NTR-GFP and nlsEos. We posit that these more mature nlsEos cells serve as hair cell progenitors. Consistent with this idea, Mtz treatment at 5 dpf that spares nlsEos cells not expressing NTR-GFP has only a small effect on hair cell regeneration while Mtz treatment at both 3 and 5 dpf results in substantial reduction in hair cell regeneration.

**Figure 12.**
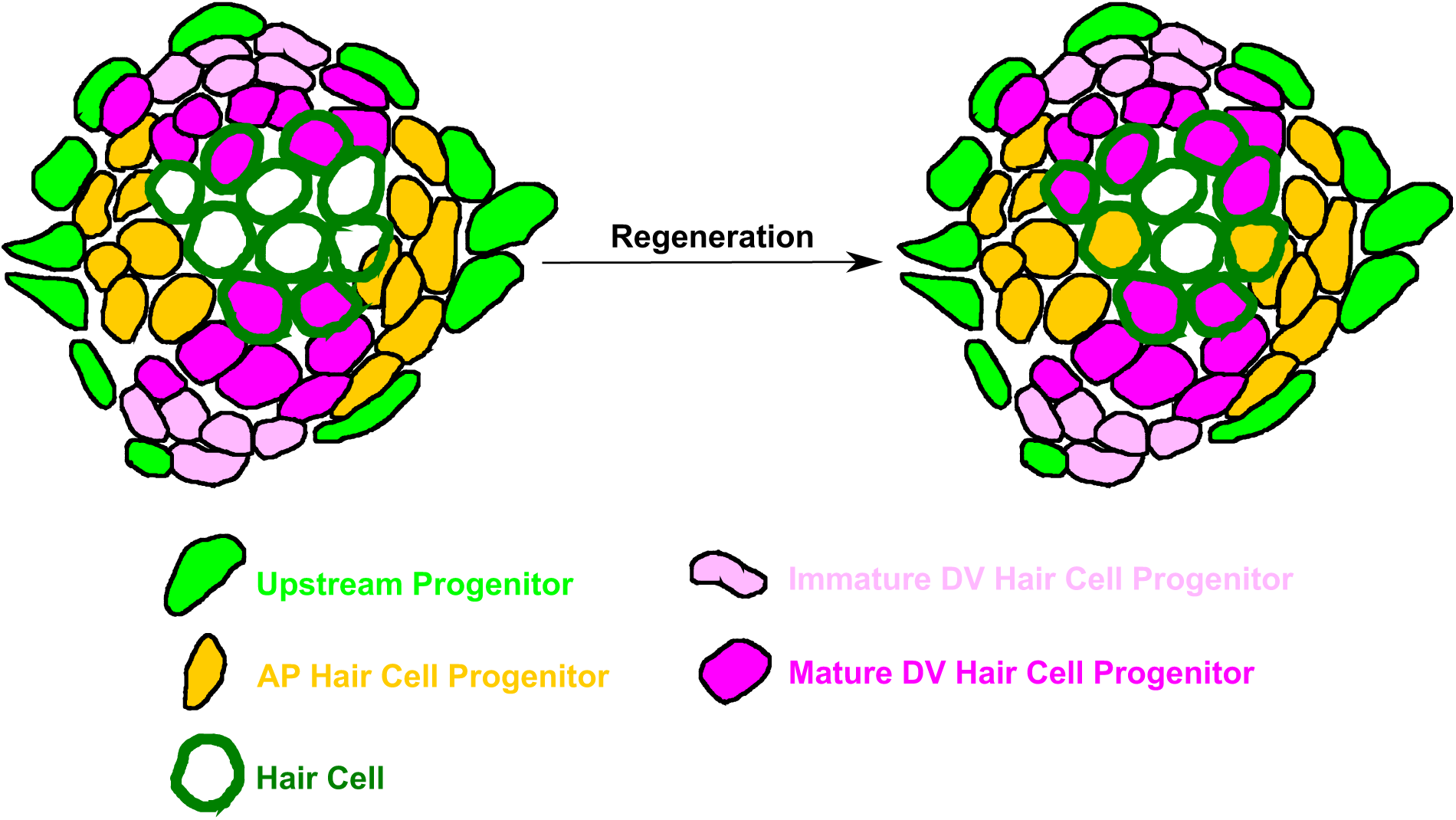
Model of neuromast progenitor identity. *Sost*:nlsEos-positive cells, located in the dorsoventral (DV) region of the neuromast, contain immature hair cell progenitors (shown in light pink) and mature hair cell progenitors (shown in magenta). Immature hair cell progenitors do not directly generate new hair cells (outlined in dark green) during regeneration, but do become mature hair cell progenitors, which comprise the majority of hair cell progenitors (see magenta-filled hair cells following regeneration). *Tnfsf10l3*:nlsEos-positive cells (shown in gold), located in the anteroposterior (AP) region of the neuromast, also serve as hair cell progenitors (see gold-filled hair cells following regeneration). Both of these populations are regulated by Notch signaling, and both can replenish immature hair cell progenitors. Finally, *sfrp1a*:nlsEos-positive cells (shown in light green), located in the periphery, do not serve as hair cell progenitors, nor are they regulated by Notch signaling. However, they are capable of replenishing immature hair cell progenitors, and can thus be classified as an upstream progenitor.

While we have identified distinct hair cell progenitor populations (DV and AP), these populations do not account for all of the hair cell progenitors in the neuromast. The combination of these cells generated 88% of new hair cells, meaning that the remaining 12% were derived from other sources. Furthermore, the AP population only accounted for 40% of new hair cells generated after neomycin treatment following DV cell ablation. Thus, there must be some other population (or populations) of support cells that are serving as hair cell progenitors that we have not labeled with our transgenic techniques. The best candidates for this role are centrally located support cells found ventral to hair cells, although the identity of these cells remains to be determined.

### The Role of Planar Cell Polarity and Progenitor Localization

Neuromasts located on the trunk develop at different times from different migrating primordia. Within a given neuromast, hair cells are arranged such that their apical stereocilia respond to directional deflection in one of two directions along the body axis. Hair cells derived from the first primordium (primI) respond along the anteroposterior axis, and hair cells derived from the second primordium (primII) respond along the dorsoventral axis (López-Schier et al. 2004; López-Schier and Hudspeth 2006). Spatial restriction of support cell proliferation is orthogonal to hair cell planar polarity, with proliferation occurring dorsoventrally in primI-derived neuromasts and anteroposteriorly in primII-derived neuromasts (Romero-Carvajal et al. 2015). This 90-degree switch between prim1- and primII-derived neuromasts is reflected in the distribution of labeled cell populations as well: *tnfsf10l3*:nlsEos and *sost*:nlsEos retain their asymmetric localization in primII-derived neuromasts, but *tnfsf10l3*:nlsEos is predominantly expressed in the dorsoventral region and *sost*:nlsEos is restricted to the anteroposterior region (data not shown). *Sfrp1a*:nlsEos expression remains limited to the periphery. Thus, within a given neuromast along the trunk, expression of *sost*:nlsEos is orthogonal to hair cell planar polarity, and that of *tnfsf10l3*:nlsEos is parallel to hair cell planar polarity.

The relationship between asymmetric progenitor localization and hair cell planar polarity remains unknown. Planar cell polarity (PCP) signaling often drives asymmetry in other tissues and has been implicated in the planar polarity of lateral line hair cells. Zebrafish deficient in Vangl2, a critical component of the PCP pathway, still develop neuromasts, but their hair cells are oriented randomly toward one another and do not respond along a single axis (López-Schier and Hudspeth 2006). Furthermore, this random orientation stems from misaligned divisions of hair cell progenitors (Mirkovic, Pylawka, and Hudspeth 2012). The transcription factor Emx2 has also been recently implicated in determining hair cell planar polarity (Jiang, Kindt, and Wu 2017). Whether these genes or other components of PCP signaling mediate the asymmetric expression of *sost*:nlsEos and *tnfsf10l3*:nlsEos remains to be determined. It would also be interesting to examine whether hair cell planar polarity is influenced by the asymmetric localization of these progenitor populations, or vice versa.

### Regeneration of Support Cells in the Absence of Hair Cell Damage

Since zebrafish are able to properly regenerate their hair cells after multiple successive insults (Cruz et al. 2015; Pinto-Teixeira et al. 2015), and both daughters of progenitors give rise to hair cells, there must be a means of replenishing hair cell progenitors. Our EdU/BrdU double labeling experiment qualitatively demonstrated that hair cell progenitors could be replenished via proliferation of other support cells. It was thus unsurprising that DV cells could themselves regenerate. It is notable, however, that DV cells could be replenished even in the absence of hair cell damage, which means that hair cell death is not the sole signal that triggers support cell proliferation. Support cell regeneration in the absence of hair cell death has been observed in mammals, as certain types of cochlear support cells (inner border cells and inner phalangeal cells) are capable of regeneration following selective ablation, a process that occurs via transdifferentiation (Mellado Lagarde et al. 2014). In contrast, zebrafish DV cell regeneration primarily occurs via proliferation, as a majority of new DV cells were EdU-positive following Mtz-induced ablation. DV cells that were not EdU-positive could have arisen after the EdU pulse, been retained due to incomplete ablation, or potentially resulted from transdifferentiation from another source.

All three labeled support cell populations were able to replenish DV cells following Mtz-induced ablation. DV cells themselves contributed nearly 60% of new DV cells, although we cannot rule out that this number is inflated due to incomplete ablation. This result suggests that DV cells choose to either generate new hair cells or replenish lost DV cells, undergoing a form of self-renewal that does not require asymmetric division. The AP population is also capable of replenishing DV cells. That both of these populations can generate new DV cells is consistent with recent findings from Viader-Llargués et al., (2018). This study defined support cells as a peripheral mantle population and a central sustentacular population. Following laser ablation of large portions of the neuromast, they found that sustentacular cells were able to regenerate mantle cells and other sustentacular cells, as well as hair cells, and could thus be considered tripotent progenitors. The transgenic lines they used to label sustentacular cells were broadly expressed, and thus should encompass both AP and DV populations. While we have not been able to selectively ablate the Peripheral cell population, and therefore cannot test whether DV and AP populations can generate them, the DV and AP populations can both generate new hair cells and new DV cells, indicating that they are both at least bipotent.

We found that the Peripheral population could also generate new DV cells following Mtz-induced ablation. Furthermore, it contributes more to DV regeneration than does the AP population. This was especially surprising since proliferation has rarely been observed in peripheral cells, at least during normal hair cell regeneration (Ma, Rubel, and Raible 2008; Romero-Carvajal et al. 2015). However, the loss of a progenitor population could be considered to be a case of extreme damage to the neuromast, thus prompting the Peripheral cell population to respond. That Peripheral cells can serve as progenitors only in extreme circumstances is consistent with the findings of Romero-Carvajal et al., 2015 and Viader-Llargués et al., 2018. Both studies suggested that mantle cells are capable of regenerating other cell types in the neuromast following extreme damage. However, the latter study found that mantle cells could only generate other mantle cells. Whether the Peripheral cell population in particular is capable of doing the same remains to be tested. Mantle cells have also been shown to proliferate following tail amputation, forming a migratory placode that forms new neuromasts along the regenerated tail (Jones and Corwin 1993; Dufourcq et al. 2006). Given the differences across studies, we have hesitated to designate the *sfrp1a*:nlsEos labeled cells as mantle cells and have instead adopted the “Peripheral” label. Since Peripheral cells can generate hair cell progenitors, we have characterized them as “upstream progenitors” (Fig. 12).

The zebrafish lateral line system continues to grow through larval and adult stages (Nuñez et al. 2009; Ledent 2002; Sapède et al. 2002), with new neuromasts formed from budding from extant neuromasts (Nuñez et al. 2009; Wada, Dambly-Chaudière, et al. 2013; Wada, Ghysen, et al. 2013) and generated anew from interneuromast cells, latent precursors deposited between neuromasts by the migrating primordium (Nuñez et al. 2009; Grant, Raible, and Piotrowski 2005). We note that the *sfrp1a*:nlsEos transgene is expressed in interneuromast cells as well as Peripheral neuromast cells. Whether these cells share similar properties to generate new neuromasts remains to be tested.

Our model of neuromast progenitor identity does bear some similarities with other regenerative tissues. Both the hair follicle and intestinal epithelium contain a niche of stem cells (bulge cells and crypt cells, respectively) which generate transit-amplifying cells that are able to generate other cell types (Ito et al. 2005; Taylor et al. 2000; Barker et al. 2007). Due to their high rate of proliferation and multipotency, the DV cells in the neuromast could be likened to these transit-amplifying cells. However, progenitors in the neuromast may bear the most similarity to those of the olfactory epithelium, which contains two distinct progenitor populations: globose basal cells (GBCs), which are transit-amplifying cells that can restore lost olfactory neurons; and horizontal basal cells (HBCs), a quiescent population that can generate multiple cell types, including GBCs, in instances of extreme damage (Iwai et al. 2008; Leung, Coulombe, and Reed 2007). In our model, the DV and Peripheral cells are comparable to the GBCs and HBCs, respectively. However, we cannot make the claim that the Peripheral population is a resident stem cell population (like bulge cells, crypt cells, and HBCs), as we do not yet know if it is capable of self-renewal or of generating every cell type within the neuromast.

### Notch Signaling Differentially Regulates Support Cell Populations

Inhibition of Notch signaling during hair cell regeneration significantly increased the number of hair cells derived from both DV and AP cells, which was not unexpected given that both are hair cell progenitor populations. However, Notch inhibition had a greater impact on the AP population than on the DV population, suggesting that it may be more strongly regulated by Notch signaling. The receptor *notch3*, in particular, is most strongly expressed in the anteroposterior portions of the neuromast (Wibowo et al. 2011; Romero-Carvajal et al. 2015). Furthermore, a transgenic reporter of Notch activity is also expressed in the anteroposterior region (Romero-Carvajal et al. 2015; Wibowo et al. 2011). It is thus likely that asymmetrically-localized Notch signaling maintains quiescence among AP cells during homeostasis and is responsible for suppressing the contribution of the AP population to hair cell regeneration (compared to DV contribution). Since Notch signaling is not as strong in the dorsoventral regions of the neuromast, the DV population is less affected and could already be more “primed” to serve as hair cell progenitors than the AP population.

Notch inhibition did not have any impact on the Peripheral population’s contribution to hair cell regeneration, indicating that Notch signaling does not suppress hair cell production by Peripheral cells. While it is difficult to tell from *in situ* expression, it seems that the Notch reporter is not active in the peripheral mantle cells (Wibowo et al. 2011; Romero-Carvajal et al. 2015). Thus, there must be some other mechanism, either intrinsic or extrinsic, that maintains relative quiescence among the Peripheral population.

It is not clear why these distinct populations of progenitors exist, as there is no clear difference in the types of hair cells they produce. Hair cells polarized in opposing direction are daughters of the final division of the hair cell progenitor (López-Schier and Hudspeth 2006). Heterogeneity has also been recently described in the synaptic responses of lateral line hair cells (Zhang et al. 2018), but these differences appear lineage-independent. Instead, the allocation of distinct progenitors may serve an advantage with respect to their differential regulation. For example, our data suggest that DV cells contribute more to homeostatic addition of new hair cells to the neuromast in the absence of damage, while both AP and DV cells are engaged after hair cell damage. AP cells were unable to overcome the loss of the DV population, suggesting that feedback mechanisms regulating both hair cell production and progenitor replacement operate independently. While both AP and DV populations are regulated by Notch signaling, this regulation appears to operate independently as Notch inhibition does not compensate for DV ablation. Independent progenitors may offer the flexibility to add new hair cells under a variety of conditions for both ongoing hair cell turnover and in the face of catastrophic hair cell loss.

## MATERIALS AND METHODS

### Fish Maintenance

Experiments were conducted on 5-8 dpf larval zebrafish (except for the double hair cell ablation experiment, which was conducted on 15-18 dpf fish). Larvae were raised in E3 embryo medium (14.97 mM NaCl, 500 μM KCL, 42 μM Na_2_HPO_4_, 150 μM KH_2_PO_4_, 1 mM CaCl_2_ dihydrate, 1 mM MgSO_4_, 0.714 mM NaHCO_3_, pH 7.2) at 28.5°C. All wildtype animals were of the AB strain. Zebrafish experiments and husbandry followed standard protocols in accordance with University of Washington Institutional Animal Care and Use Committee guidelines.

### Plasmid Construction

The myo6b:mKate2 construct was generated via the Gateway Tol2 system (Invitrogen). A pME-mKate2 (the mKate2 sequence being cloned from pMTB-Multibow-mfR, Addgene #60991) construct was generated via BP Recombination, and then a pDEST-myo6b:mKate2 construct was generated via LR recombination of p5E-myo6b, pME-mKate2, p3E-pA, and pDEST-iTol2-pA2 vectors. The mbait-GFP construct was a gift from Shin-Ichi Higashijima’s lab. The mbait-nlsEos construct was also generated via Gateway LR recombination of p5E-mbait/HSP70l, pME-nlsEos, p3E-pA, and pDEST-iTol2-pA2 vectors. The mbait-epNTR-GFP construct was generated via Gibson assembly, inserting the coding sequence of epNTR (cloned from pCS2-epNTR obtained from Harold Burgess’ lab) plus a small linker sequence in front of the GFP in the original pBSK mbait-GFP vector. All plasmids were maxi prepped (Qiagen) prior to injection.

### CRISPR Guides

Gene-specific guide RNA (gRNA) sequences were as follows:

*sfrp1a*: GTCTGGCCTAAAGAGACCAG

*tnfsf10l3*: GGGCTTGTATAGGAGTCACG

*sost*: GGGAGTGAGCAGGGATGCAA

GGGCGAAGAACGGTGGAAGG

All gRNAs were synthesized according to the protocol outlined in (Shah et al. 2015), but were purified using a Zymo RNA Clean & Concentrator kit. Upon purification, gRNAs were diluted to 1 μg/μL, aliquoted into 4 μL aliquots, and stored at -80°C. *Sfrp1a* and *tnfsf10l3* gRNA sequences were designed via http://crispr.mit.edu, and the *sost* gRNA sequences were designed via http://crisprscan.org. The *tnfsf10l3* gRNA was targeted upstream of the gene’s start ATG codon, whereas the *sfrp1a* and *sost* guides were targeted to exons.

### Tol2 Transgenesis

The Tg[myo6b:mKate2]^w218^ (hereafter called myo6:mKate2) line was generated via Tol2 transgenesis. 1-2 nL of a 5 μL injection mix consisting of 20 ng/μL myo6b:mKate2 plasmid, 40 ng/μL transposase mRNA, and 0.2% phenol red were injected into single cell wildtype embryos. Larvae were screened for expression at 3 dpf and transgenic F_0_ larvae were grown to adulthood. F_0_ adults were outcrossed to wildtype fish, transgenic offspring were once again grown to adulthood, and the resulting adults were used to maintain a stable line.

### CRISPR-mediated Transgenesis

All support cell transgenic lines were generated via CRISPR-mediated transgenesis. For most injections, a 5 μL injection mix was made consisting of 200 ng/μL gene-specific gRNA, 200 ng/μL mbait gRNA, 800 ng/μL Cas9 protein (PNA Bio #CP02), 20 ng/μL mbait-reporter plasmid, and 0.24% phenol red. The gRNAs and Cas9 protein were mixed together first, then heated at 37°C for 10 minutes, after which the other components were added. In the case of *sost*, in which two gRNAs were co-injected, each gRNA was added to the mix at a final concentration of 100 ng/μL (so 200 ng/μL of total guide-specific gRNA). When reconstituting the Cas9 protein, DTT was added to a final concentration of 1 mM DTT. This is highly recommended to reduce needle clogging during the injection process!! 1-2 nL of these injection mixes were injected into single cell wildtype embryos. Larvae were screened for expression at 3 dpf and transgenic F_0_ larvae were grown to adulthood. F_0_ adults were outcrossed to wildtype fish, transgenic offspring were once again grown to adulthood, and the resulting adults were used to maintain a stable line.

### Photoconversion

In order to photoconvert multiple nlsEos fish at once, larvae were transferred to a 60 ×15mm petri dish and placed in a freezer box lined with aluminum foil. Then, an iLumen 8 UV flashlight (procured from Amazon) was placed over the dish and turned on for 15 minutes. Following the UV pulse, larvae were returned to standard petri dishes to await experimentation.

### Drug Treatments

For all drug treatments, zebrafish larvae were placed in baskets in 6 well plates to facilitate transfer of larvae between media. Larvae were treated at 5 dpf unless otherwise noted. All wells contained 10 mL of drug, E3 embryo medium with the same effective % DMSO as the drug (for mock treatments), or plain E3 embryo medium for washout. Following treatment, the fish were washed twice into fresh E3 embryo medium by moving the baskets into adjacent wells in the row, then washed a third time by transferring them into a 100 mm petri dish with fresh E3 medium. All drugs were diluted in E3 embryo medium. The drug treatment paradigms were as follows: for hair cell ablation, 400 μM neomycin (Sigma) for 30 minutes; for *sost*:NTR ablation, 10 mM metronidazole (Mtz; Sigma) with 1% DMSO; for Notch inhibition, 50 μM LY411575 (LY; Sigma) for 24 hours; for the *sost* ablation/Notch inhibition experiment (Fig. 10): 10mM Mtz/50 μM LY for 8 hours, then 50 μM LY for 16 hours.

For double hair cell ablation studies, larvae that were treated with neomycin were raised on a nursery in the UW fish facility beginning at 7 dpf and then treated with 400 μM neomycin again at 15 dpf in standard petri dishes (10 days following the first neomycin treatment). These juvenile fish were washed into fresh system water multiple times before being returned to the nursery and were then fixed three days later (18 dpf).

### EdU and BrdU Treatments

Following hair cell ablation with neomycin, larvae were incubated in 500 μM F-ara-EdU (Sigma #T511293) for 24 hours. Following *sost*:NTR ablation with Mtz, larvae were incubated in the same concentration of EdU for 48 hours. Larvae were placed into fresh EdU after the first 24 hours. F-ara-EdU was originally reconstituted in 50% H_2_O and 50% DMSO to 50 mM, so the working concentration of DMSO of 500 μM was 0.5%. In the case of the double ablation studies, juvenile fish were incubated in 10mM BrdU (Sigma) with 1% DMSO in system water (used in the UW fish facility) following the second neomycin treatment for 24 hours. Following treatment, larvae were washed in fresh system water several times.

### Immunohistochemistry

Zebrafish larvae were fixed in 4% paraformaldehyde in PBS containing 4% sucrose for either 2 hours at room temperature or overnight at 4°C. Larvae were then washed 3 times (20 minutes each) in PBS containing 0.1% Tween20 (PBT), incubated for 30 minutes in distilled water, then incubated in antibody block (5% heat-inactivated goat serum in PBS containing 0.2% Triton, 1% DMSO, 0.02% sodium azide, and 0.2% BSA) for at least one hour at room temperature. Larvae were then incubated in mouse anti-parvalbumin or rabbit anti-GFP (or sometimes both simultaneously) diluted 1:500 in antibody block overnight at 4°C. The next day, larvae were once again washed 3 times (20 minutes each) in PBT, then incubated in a fluorescently-conjugated secondary antibody (Invitrogen, Alexa Fluor 488, 568, and/or 647) diluted 1:1000 in antibody block for 4-5 hours at room temperature. From this point onward, larvae were protected from light. Larvae were then rinsed 3 times (10 minutes each) in PBT and then stored in antibody block at 4°C until imaging. For BrdU immunohistochemistry, juvenile fish were rinsed once in 1N HCl, then incubated in 1N HCl following washout of Click-iT reaction mix (see below). IHC proceeded as above, except that the antibody block contained 10% goat serum and the fish were incubated in mouse anti-BrdU at a dilution of 1:100. All wash and incubation steps occurred with rocking.

### Click-iT

Cells that had incorporated F-ara-EdU were visualized via a Click-iT reaction. In the case of the double hair cell ablation experiment, Click-iT was performed before immunohistochemistry. Following fixation, fish were washed 3 times (10 minutes each) in PBT, then permeabilized in PBS containing 0.5% Triton-X for 30 minutes, then washed another 3 times (10 minutes each) in PBS. Next, fish were incubated for 1 hour at room temperature in a Click-iT reaction mix consisting of 2mM CuSO_4_, 10 μM Alexa Fluor 555 azide, and 20 mM sodium ascorbate in PBS (made fresh). Fish were protected from light from this point onwards. Afterwards, the standard IHC protocol listed above was performed (beginning with the 3 20-minute washes in PBT). For the *sost*:NTR regeneration experiment, the Click-iT reaction was performed after IHC. Following incubation in secondary antibody, larvae were washed 3 times (10 minutes each) in PBS, then incubated in 700 μL of the Click-iT reaction mix (again, made immediately prior to incubation) for 1 hour at room temperature. Larvae were then washed 6 times (20 minutes each) in PBT to ensure proper clearing of background labeling and stored in antibody block at 4°C until imaging.

### Confocal Imaging

With the exception of imaging requiring a far-red laser (Fig. 1F-N, Fig. 3C-I, Fig. 5), all imaging was performed using an inverted Marianas spinning disk system (Intelligent Imaging Innovations, 3i) with an Evolve 10 MHz EMCCD camera (Photometrics) and a Zeiss C-Apochromat 63x/1.2W numerical aperture water objective. All imaging experiments were conducted with fixed larvae ages 5-8 dpf. Fish were placed in a chambered borosilicate coverglass (Lab-Tek) containing 2.5-2.5 mL E3 embryo medium and oriented on their sides with a slice anchor harp (Harvard Instruments). Imaging was performed at ambient temperature, generally 25°C. Fish were positioned on their sides against the cover glass in order to image the first 5 primary neuromasts of the posterior lateral line (P1-P5). All imaging was performed with an intensification of 650, a gain of 3, an exposure time between 25-1500 ms (depending on the brightness of the signal) for 488, 561, and 405 lasers, and a step size of 1 μm. All 3i Slidebook images were exported as.tif files to Fiji.

In cases when a far-red laser was required, imaging was performed on a Zeiss LSM 880 microscope with a Zeiss C-Apochromat 40x/1.2W numerical water objective. Fish were immersed in a solution of 50%glycerol/50% PBS, and then mounted on a plain microscope slide (Richard-Allen) beneath a triple wholemount coverslip. Imaging was performed at ambient temperature, generally 25°C. Fish were positioned on their sides against the cover glass in order to image the first 5 primary neuromasts of the posterior lateral line (P1-P5). For the double hair cell ablation experiment, following fixation the tails of fish were cut off and mounted underneath a single wholemount coverslip. The 3 neuromasts of the terminal cluster were imaged per tail. All imaging was performed at 4-5x digital zoom with master gain between 500-800 for 488, 561, and 647 lasers, and a step size of 1 μm. All images were captured through the Zen Black software and opened in Fiji as.czi files.

### Statistical Analysis

All statistical analyses were done with GraphPad Prism 6.0. The Mann Whitney U test was used for comparisons between 2 groups, whereas the Kruskal-Wallis test, with a Dunn’s post-test, was used for comparisons between 3 or more groups. Statistical significance was set at p = 0.05.

## Acknowledgements

We would like to thank David White and the staff of the UW Fish Facility for animal care and maintenance, Ivan Cruz for providing the *sost* gRNA, Shin-Ichi Higashijima for the mbait-GFP vector, Harold Burgess for the pCS2-epNTR vector, Madeleine Hewitt for assistance with data analysis, and Sarah Pickett for critical reading of the manuscript.

## AUTHOR CONTRIBUTIONS

Conceptualization and Methodology: E.D.T. and D.W.R.; Investigation: E.D.T.; Writing – Original Draft: E.D.T.; Writing – Review & Editing: E.D.T. and D.W.R.; Funding Acquisition: D.W.R.; Supervision: D.W.R.

---

**Figure 2 – figure supplement 1.**
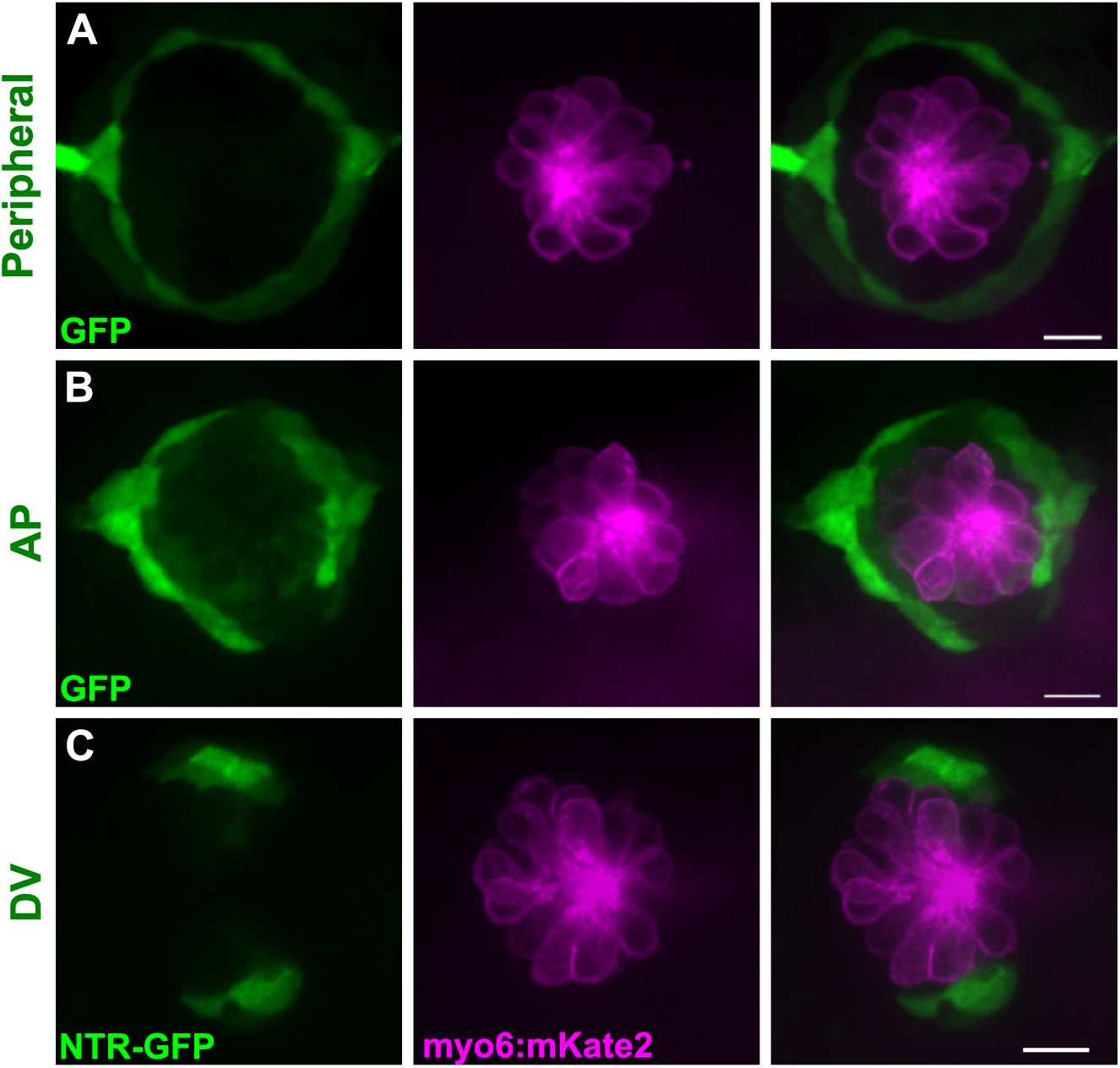
Support cell transgenes are not expressed in hair cells. **(A-C)** Maximum projections of neuromasts from Tg[*sfrp1a*:GFP]^w222^ (Peripheral, A), Tg[*tnfsf10l3*:GFP]^w223^ (AV, B), and *sost*:NTR-GFP (DV, C) fish. GFP-positive cells are shown in green, and hair cells are shown in magenta via myo6:mKate2. In all three populations, there is no GFP expression in hair cells. Scale bar = 10 μm.

**Figure 6 – figure supplement 1.**
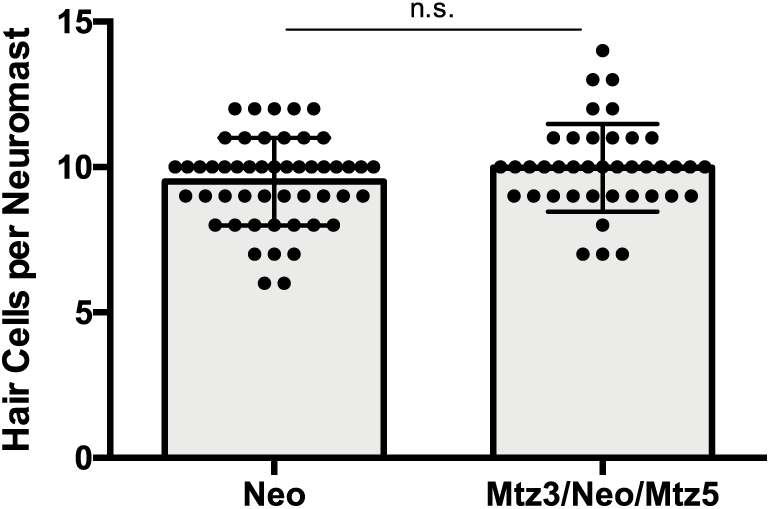
Mtz treatment does not inherently impact hair cell regeneration. Total number of hair cells per neuromast following regular hair cell regeneration (Neo) or DV cell-ablated regeneration (Mtz3/Neo/Mtz5) in non-transgenic siblings of *sost*:NTR-GFP fish. Neo: 9.5 ± 1.50, n = 50 neuromasts; Mtz3/Neo/Mtz5: 9.98 ± 1.51, n = 40 neuromasts; mean ± SD; Mann Whitney U test, p = 0.2317.

## REFERENCES

Barker, Nick, Johan H. van Es, Jeroen Kuipers, Pekka Kujala, Maaike van den Born, Miranda Cozijnsen, Andrea Haegebarth, et al. 2007. “Identification of Stem Cells in Small Intestine and Colon by Marker Gene Lgr5.” Nature 449 (7165): 1003–7. DOI:10.1038/nature06196.

Choi, Rhea, and Bradley J. Goldstein. 2018. “Olfactory Epithelium: Cells, Clinical Disorders, and Insights from an Adult Stem Cell Niche.” Laryngoscope Investigative Otolaryngology 3 (1). Wiley-Blackwell: 35–42. DOI:10.1002/lio2.135.

Cruz, Ivan A., Ryan Kappedal, Scott M. Mackenzie, Dale W. Hailey, Trevor L. Hoffman, Thomas F. Schilling, and David W. Raible. 2015. “Robust Regeneration of Adult Zebrafish Lateral Line Hair Cells Reflects Continued Precursor Pool Maintenance.” Developmental Biology 402 (2). Academic Press Inc.: 229–38. DOI:10.1016/j.ydbio.2015.03.019.

Curado, Silvia, Ryan M. Anderson, Benno Jungblut, Jeff Mumm, Eric Schroeter, and Didier Y.R. Stainier. 2007. “Conditional Targeted Cell Ablation in Zebrafish: A New Tool for Regeneration Studies.” Developmental Dynamics 236 (4): 1025–35. DOI:10.1002/dvdy.21100.

Dufourcq, Pascale, Myriam Roussigné, Patrick Blader, Frédéric Rosa, Nadine Peyrieras, and Sophie Vriz. 2006. “Mechano-Sensory Organ Regeneration in Adults: The Zebrafish Lateral Line as a Model.” Molecular and Cellular Neuroscience 33 (2). Academic Press: 180–87. DOI:10.1016/J.MCN.2006.07.005.

Grant, Kelly a., David W. Raible, and Tatjana Piotrowski. 2005. “Regulation of Latent Sensory Hair Cell Precursors by Glia in the Zebrafish Lateral Line.” Neuron 45 (1): 69–80. DOI:10.1016/j.neuron.2004.12.020.

Harris, Julie A, Alan G Cheng, Lisa L Cunningham, Glen MacDonald, David W Raible, and Edwin W Rubel. 2003. “Neomycin-Induced Hair Cell Death and Rapid Regeneration in the Lateral Line of Zebrafish (Danio Rerio).” Journal of the Association for Research in Otolaryngology?: JARO 4 (2): 219–34. DOI:10.1007/s10162-002-3022-x.

Hsu, Ya-Chieh, Lishi Li, and Elaine Fuchs. 2014. “Emerging Interactions between Skin Stem Cells and Their Niches.” Nature Medicine 20 (8): 847–56. DOI:10.1038/nm.3643.

Ito, Mayumi, Yaping Liu, Zaixin Yang, Jane Nguyen, Fan Liang, Rebecca J Morris, and George Cotsarelis. 2005. “Stem Cells in the Hair Follicle Bulge Contribute to Wound Repair but Not to Homeostasis of the Epidermis.” Nature Medicine 11 (12): 1351–54. DOI:10.1038/nm1328.

Iwai, Naomi, Zhijian Zhou, Dennis R Roop, and Richard R Behringer. 2008. “Horizontal Basal Cells Are Multipotent Progenitors in Normal and Injured Adult Olfactory Epithelium.” Stem Cells (Dayton, Ohio) 26 (5). NIH Public Access: 1298–1306. DOI:10.1634/stemcells.2007-0891.

Jiang, Tao, Katie Kindt, and Doris K Wu. 2017. “Transcription Factor Emx2 Controls Stereociliary Bundle Orientation of Sensory Hair Cells.” ELife 6. eLife Sciences Publications, Ltd. DOI:10.7554/eLife.23661.

Jones, JE, and JT Corwin. 1993. “Replacement of Lateral Line Sensory Organs during Tail Regeneration in Salamanders: Identification of Progenitor Cells and Analysis of Leukocyte Activity.” Journal of Neuroscience 13 (3). Society for Neuroscience: 1022–34. DOI:10.1523/JNEUROSCI.13-03-01022.1993.

Kimura, Yukiko, Yu Hisano, Atsuo Kawahara, and Shin-Ichi Higashijima. 2014. “Efficient Generation of Knock-in Transgenic Zebrafish Carrying Reporter/Driver Genes by CRISPR/Cas9-Mediated Genome Engineering.” Scientific Reports. DOI:10.1038/srep06545.

Ledent, Valérie. 2002. “Postembryonic Development of the Posterior Lateral Line in Zebrafish.” Development 129 (3).

Leung, Cheuk T, Pierre A Coulombe, and Randall R Reed. 2007. “Contribution of Olfactory Neural Stem Cells to Tissue Maintenance and Regeneration.” Nature Neuroscience 10 (6): 720–26. DOI:10.1038/nn1882.

López-Schier, Hernán, and A J Hudspeth. 2006. “A Two-Step Mechanism Underlies the Planar Polarization of Regenerating Sensory Hair Cells.” Proceedings of the National Academy of Sciences of the United States of America 103 (49): 18615–20. DOI:10.1073/pnas.0608536103.

López-Schier, Hernán, Catherine J. Starr, James a. Kappler, Richard Kollmar, and a. J. Hudspeth. 2004. “Directional Cell Migration Establishes the Axes of Planar Polarity in the Posterior Lateral-Line Organ of the Zebrafish.” Developmental Cell 7 (3): 401–12. DOI:10.1016/j.devcel.2004.07.018.

Ma, Eva Y., Edwin W. Rubel, and David W. Raible. 2008. “Notch Signaling Regulates the Extent of Hair Cell Regeneration in the Zebrafish Lateral Line.” The Journal of Neuroscience 28 (9): 2261–73. DOI:10.1523/JNEUROSCI.4372-07.2008.

Mackenzie, Scott M., and David W. Raible. 2012. “Proliferative Regeneration of Zebrafish Lateral Line Hair Cells after Different Ototoxic Insults.” PLoS ONE 7 (10): 1–8. DOI:10.1371/journal.pone.0047257.

McMenamin, Sarah K., Emily J. Bain, Anna E. McCann, Larissa B. Patterson, Dae Seok Eom, Zachary P. Waller, James C. Hamill, Julie A. Kuhlman, Judith S. Eisen, and David M. Parichy. 2014. “Thyroid Hormone-Dependent Adult Pigment Cell Lineage and Pattern in Zebrafish.” Science (New York, N.Y.) 345 (6202). NIH Public Access: 1358. DOI:10.1126/SCIENCE.1256251.

Mellado Lagarde, Marcia M, Guoqiang Wan, LingLi Zhang, Angelica R Gigliello, John J McInnis, Yingxin Zhang, Dwight Bergles, Jian Zuo, and Gabriel Corfas. 2014. “Spontaneous Regeneration of Cochlear Supporting Cells after Neonatal Ablation Ensures Hearing in the Adult Mouse.” Proceedings of the National Academy of Sciences of the United States of America 111 (47). National Academy of Sciences: 16919–24. DOI:10.1073/pnas.1408064111.

Mirkovic, I., S. Pylawka, and a. J. Hudspeth. 2012. “Rearrangements between Differentiating Hair Cells Coordinate Planar Polarity and the Establishment of Mirror Symmetry in Lateral-Line Neuromasts.” Biology Open 1 (5): 498–505. DOI:10.1242/bio.2012570.

Mizutari, Kunio, Masato Fujioka, Makoto Hosoya, Naomi Bramhall, Hideyuki Hirotaka James Okano, Hideyuki Hirotaka James Okano, and Albert S B Edge. 2013. “Notch Inhibition Induces Cochlear Hair Cell Regeneration and Recovery of Hearing after Acoustic Trauma.” Neuron 77 (1). Elsevier Inc.: 58–69. DOI:10.1016/j.neuron.2012.10.032.

Neef, A. B., and N. W. Luedtke. 2011. “Dynamic Metabolic Labeling of DNA in Vivo with Arabinosyl Nucleosides.” Proceedings of the National Academy of Sciences 108 (51): 20404–9. DOI:10.1073/pnas.1101126108.

Nuñez, Viviana a., Andres F. Sarrazin, Nicolas Cubedo, Miguel L. Allende, Christine Dambly-Chaudière, and Alain Ghysen. 2009. “Postembryonic Development of the Posterior Lateral Line in the Zebrafish.” Evolution and Development 11 (4): 391–404. DOI:10.1111/j.1525-142X.2009.00346.x.

Ota, Satoshi, Kiyohito Taimatsu, Kanoko Yanagi, Tomohiro Namiki, Rie Ohga, Shin-ichi Higashijima, and Atsuo Kawahara. 2016. “Functional Visualization and Disruption of Targeted Genes Using CRISPR/Cas9-Mediated EGFP Reporter Integration in Zebrafish.” Scientific Reports 6 (1): 34991. DOI:10.1038/srep34991.

Pinto-Teixeira, F., O. Viader-Llargues, E. Torres-Mejia, M. Turan, E. Gonzalez-Gualda, L. Pola-Morell, and H. Lopez-Schier. 2015. “Inexhaustible Hair-Cell Regeneration in Young and Aged Zebrafish.” Biology Open 4 (7): 903–9. DOI:10.1242/bio.012112.

Romero-Carvajal, Andrés, Joaquín Navajas Acedo, Linjia Jiang, Agne Kozlovskaja-Gumbriene, Richard Alexander, Hua Li, Tatjana Piotrowski, et al. 2015. “Regeneration of Sensory Hair Cells Requires Localized Interactions between the Notch and Wnt Pathways.” Developmental Cell 34 (3). NIH Public Access: 267–82. DOI:10.1016/j.devcel.2015.05.025.

Rompolas, Panteleimon, and Valentina Greco. 2014. “Stem Cell Dynamics in the Hair Follicle Niche.” Seminars in Cell & Developmental Biology 25–26. NIH Public Access: 34–42. DOI:10.1016/j.semcdb.2013.12.005.

Santos, António J M, Yuan-Hung Lo, Amanda T Mah, and Calvin J Kuo. 2018. “The Intestinal Stem Cell Niche: Homeostasis and Adaptations.” Trends in Cell Biology 0 (0). Elsevier. DOI:10.1016/j.tcb.2018.08.001.

Sapède, Dora, Nicolas Gompel, Christine Dambly-Chaudière, and Alain Ghysen. 2002. “Cell Migration in the Postembryonic Development of the Fish Lateral Line.” Development (Cambridge, England) 129 (3): 605–15. http://www.ncbi.nlm.nih.gov/pubmed/11830562.

Schwob, James E, Woochan Jang, Eric H Holbrook, Brian Lin, Daniel B Herrick, Jesse N Peterson, and Julie Hewitt Coleman. 2017. “Stem and Progenitor Cells of the Mammalian Olfactory Epithelium: Taking Poietic License.” The Journal of Comparative Neurology 525 (4). NIH Public Access: 1034–54. DOI:10.1002/cne.24105.

Shah, Arish N, Crystal F Davey, Alex C Whitebirch, Adam C Miller, and Cecilia B Moens. 2015. “Rapid Reverse Genetic Screening Using CRISPR in Zebrafish.” Nature Methods 12 (6). NIH Public Access: 535–40. DOI:10.1038/nmeth.3360.

Tabor, Kathryn M, Sadie A Bergeron, Eric J Horstick, Diana C Jordan, Vilma Aho, Tarja Porkka-Heiskanen, Gal Haspel, and Harold A Burgess. 2014. “Direct Activation of the Mauthner Cell by Electric Field Pulses Drives Ultrarapid Escape Responses.” Journal of Neurophysiology 112 (4). American Physiological Society: 834–44. DOI:10.1152/jn.00228.2014.

Taylor, G, M S Lehrer, P J Jensen, T T Sun, and R M Lavker. 2000. “Involvement of Follicular Stem Cells in Forming Not Only the Follicle but Also the Epidermis.” Cell 102 (4): 451–61. http://www.ncbi.nlm.nih.gov/pubmed/10966107.

Thomas, Eric D., Ivan A. Cruz, Dale W. Hailey, and David W. Raible. 2015. “There and Back Again: Development and Regeneration of the Zebrafish Lateral Line System.” Wiley Interdisciplinary Reviews: Developmental Biology 4 (1): 1–16. DOI:10.1002/wdev.160.

Viader-Llargués, Oriol, Valerio Lupperger, Laura Pola-Morell, Carsten Marr, and Hernán López-Schier. 2018. “Live Cell-Lineage Tracing and Machine Learning Reveal Patterns of Organ Regeneration.” ELife 7 (March). DOI:10.7554/eLife.30823.

Wada, Hironori, Christine Dambly-Chaudière, Koichi Kawakami, and Alain Ghysen. 2013. “Innervation Is Required for Sense Organ Development in the Lateral Line System of Adult Zebrafish.” Proceedings of the National Academy of Sciences of the United States of America 110 (14): 5659–64. DOI:10.1073/pnas.1214004110.

Wada, Hironori, Alain Ghysen, Kazuhide Asakawa, Gembu Abe, Tohru Ishitani, and Koichi Kawakami. 2013. “Wnt/Dkk Negative Feedback Regulates Sensory Organ Size in Zebrafish.” Current Biology 23 (16). Cell Press: 1559–65. DOI:10.1016/J.CUB.2013.06.035.

Wibowo, Indra, Filipe Pinto-Teixeira, Chie Satou, Shin-ichi Higashijima, and Hernán López-Schier. 2011. “Compartmentalized Notch Signaling Sustains Epithelial Mirror Symmetry.” Development (Cambridge, England) 138 (6): 1143–52. DOI:10.1242/dev.060566.

Wiedenmann, J., S. Ivanchenko, F. Oswald, F. Schmitt, C. Rocker, A. Salih, K.-D. Spindler, and G. U. Nienhaus. 2004. “EosFP, a Fluorescent Marker Protein with UV-Inducible Green-to-Red Fluorescence Conversion.” Proceedings of the National Academy of Sciences 101 (45): 15905–10. DOI:10.1073/pnas.0403668101.

Williams, J. a., and N. Holder. 2000. “Cell Turnover in Neuromasts of Zebrafish Larvae.” Hearing Research 143 (1–2): 171–81. DOI:10.1016/S0378-5955(00)00039-3.

Xiao, Tong, Tobias Roeser, Wendy Staub, Herwig Baier, C Nüsslein-Volhard, and F Bonhoeffer. 2005. “A GFP-Based Genetic Screen Reveals Mutations That Disrupt the Architecture of the Zebrafish Retinotectal Projection.” Development (Cambridge, England) 132 (13). The Company of Biologists Ltd: 2955–67. DOI:10.1242/dev.01861.

Yousefi, Maryam, Linheng Li, and Christopher J Lengner. 2017. “Hierarchy and Plasticity in the Intestinal Stem Cell Compartment.” Trends in Cell Biology 27 (10). NIH Public Access: 753–64. DOI:10.1016/j.tcb.2017.06.006.

Zhang, Qiuxiang, Suna Li, Hiu-Tung C. Wong, Xinyi J. He, Alisha Beirl, Ronald S. Petralia, Ya-Xian Wang, and Katie S. Kindt. 2018. “Synaptically Silent Sensory Hair Cells in Zebrafish Are Recruited after Damage.” Nature Communications 9 (1). Nature Publishing Group: 1388. DOI:10.1038/s41467-018-03806-8.

